# Ancestry predicts a heritable proliferation state in *Arabidopsis*

**DOI:** 10.64898/2026.07.12.738078

**Authors:** Chunghee Lee, Soomin Lee, Donguk Gwon, Samsad Razzaque, Hyobin Jeong, Wolfgang Busch, Todd P. Michael, Sanghwa Lee

## Abstract

- Expression variation in natural plant populations is usually read gene by gene, as a response to the environment. Whether a leading axis of that variation is more strongly associated with ancestry than with home-site environment, and whether it is heritable, remain unresolved.
- We integrated transcriptomes, genotypes, environmental records and phenotypes from 665 *Arabidopsis thaliana* accessions grown under one condition.
- Genomic ancestry uniquely explained 5.0% of the dominant expression axis, compared with 2.6% for home-site environment, and exceeded environment under all four allocations of geography and the circadian covariate (1.9- to 6.2-fold). This axis was associated with a heritable cell-cycle- and ribosome-enriched module (prolifME; *h²* = 0.19), linked to four loci with widespread *trans*-expression associations and associated with smaller plants in two independent experiments and with slower growth under controlled conditions.
- The advantage is concentrated on the leading expression axes and narrows across the full transcriptome. The gene weights defining the *Arabidopsis* axis were correlated with independently derived profiles in rice and maize, with the correspondence carried mainly by photosynthesis-associated genes. Ancestry therefore predicts the leading expression axes, and the heritable proliferation module that carries the first axis tracks growth phenotypes independently of ancestry.

## Introduction

Plants occupy remarkably diverse environments, yet their core cellular programs — cell division, ribosome biogenesis and translation — are shared across genetic backgrounds and species, and are required under all conditions (Julca *et al*. 2021). Whether gene expression variation across natural plant populations is shaped primarily by the environments plants experience, or by intrinsic programs that organize transcriptome-wide variation irrespective of environmental context, is not settled.

Across biological systems, high-dimensional molecular data often resolve into dominant axes reflecting coordinated programs. A growth-versus-stress axis organizes yeast responses (Gasch *et al*. 2000), co-expression networks recover conserved developmental programs in the human brain (Oldham *et al*. 2008), and growth-associated programs structure natural transcriptome variation in *Caenorhabditis elegans* (Zhang *et al*. 2022). Plant population studies, however, have focused mainly on environmental response and local adaptation (Des Marais & Juenger 2010; Lasky *et al*. 2012; Frachon *et al*. 2017). Whether inherited programs similarly organize population-scale plant transcriptomes has not been tested.

Natural *Arabidopsis thaliana* populations vary at the molecular level along environmental gradients. Both population history and climate structure DNA methylation (Kawakatsu *et al*. 2016), and genome-wide scans detect climatic selection (Hancock *et al*. 2011; Fournier-Level *et al*. 2011; Exposito-Alonso *et al*. 2019). Yet what organizes the major axes of expression across accessions, and whether that structure is inherited or environmentally imposed, remains unknown. This differs from asking which genes respond to particular conditions: an axis may be reproducible across a species even when no single gene responds strongly to a measured variable. That ancestry predicts many *Arabidopsis* traits is not in itself surprising. Our question is whether a coherent, heritable program sits on the expression axis that ancestry predicts, and whether that program reaches the whole-plant phenotype.

A second question follows immediately. If an axis of expression variation is organized by ancestry, is it inherited in the quantitative-genetic sense, or does it merely co-vary with the leading genomic principal components used to summarize ancestry? The two are separable, because SNP heritability estimated on a genomic relationship matrix can be conditioned on those genomic components. We therefore treat the heritability of the axis, and of the covariates used to model it, as part of the result rather than as an assumption.

We therefore combined transcriptomes and genotypes with home-site climate and soil records and with archived phenotypes for 665 natural accessions (1001 Genomes Consortium 2016; Fick & Hijmans 2017; Poggio et al. 2021; Togninalli et al. 2020), and extended the analysis to rice and maize through orthogroup mapping (Emms & Kelly 2019). All accessions were grown in a single controlled condition, so the environmental predictors describe each accession’s site of origin rather than the conditions it experienced. The comparison is between ancestry-associated and home-site-environment-associated components of expression variation under a common growth regime. We show that ancestry predicts the dominant expression axis better than home-site environment. That axis corresponds to a heritable proliferation module with a bipartite motif structure and four loci showing widespread trans associations. The module is associated with growth-strategy phenotypes, and *Arabidopsis* gene weights correlate with independently derived expression profiles in rice and maize.

## Materials and Methods

### Transcriptome, genotype and environmental data

Transcriptomes for 665 natural *Arabidopsis* accessions were obtained from the 1001 Genomes RNA-sequencing dataset (GSE80744; 1001 Genomes Consortium 2016). They were supplied as upper-quartile normalized counts and z-scored per gene across accessions. One accession (ID 7427) lacks geographic coordinates and was excluded wherever coordinates enter the model, giving n = 664 for those analyses and n = 665 otherwise. Genomic principal components (genomicPC1–10) were computed from 1001 Genomes SNP data. Home-site environmental variables comprised 19 bioclimatic variables from WorldClim v2.1 (Fick & Hijmans 2017) and eight soil properties from SoilGrids at 0–5 cm depth (Poggio et al. 2021; Methods S1). The bioclimatic predictors are strongly collinear, with 17 of the 19 exceeding a variance inflation factor of 10 (Fig. 1a; Fig. S1a), so the environmental block was summarized by principal components. Phenotypes were taken from AraPheno (Togninalli et al. 2020). Two were measured under controlled conditions (Vasseur et al. 2018): rosette dry mass (ID 390) and growth rate (ID 391). A third, plant size in a Swedish common garden, provides an environmentally controlled growth measure (ID 115, n = 80; Ågren & Schemske 2012).

**Fig. 1.**
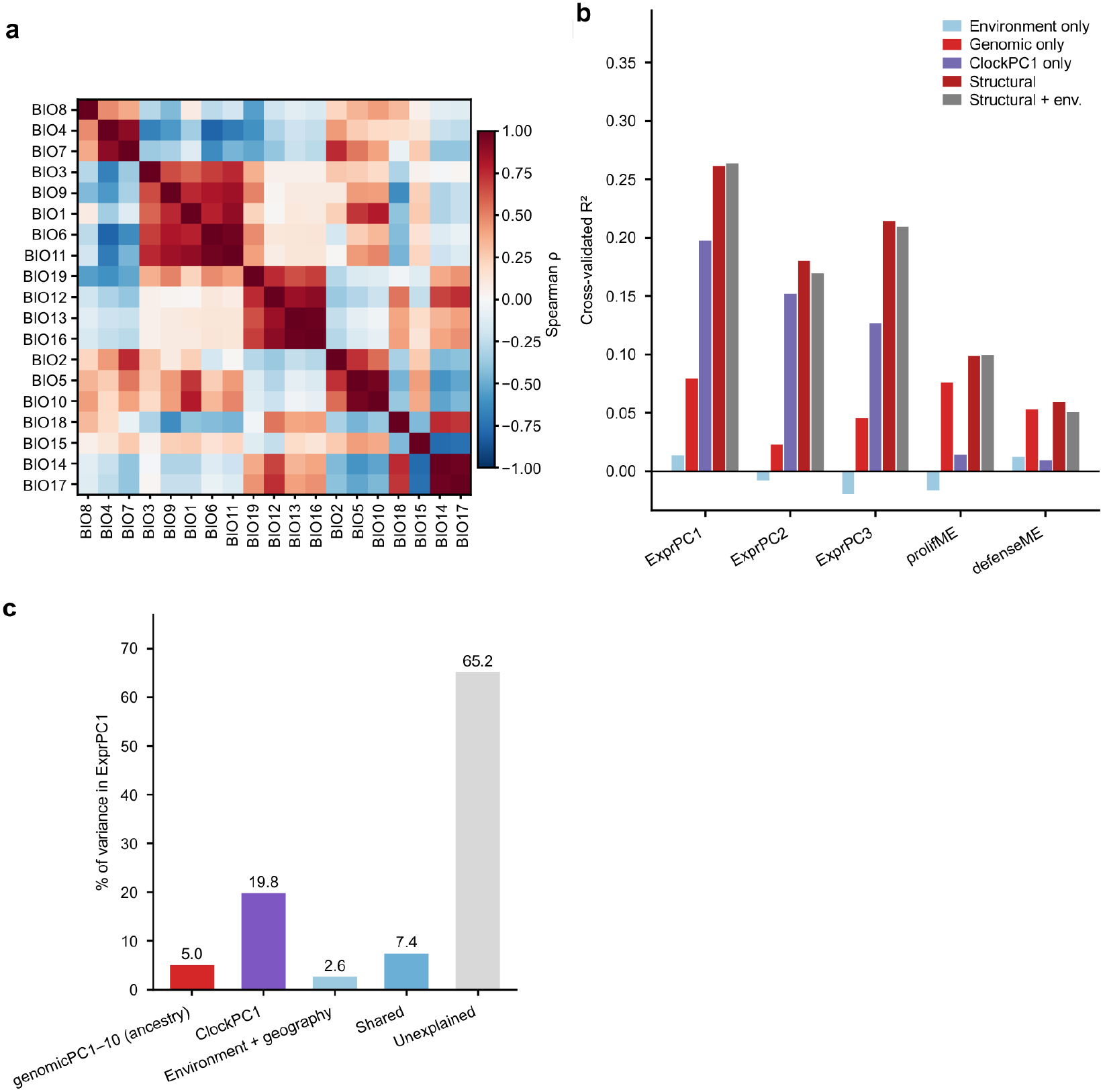
Structural predictors outpredict home-site environment on the dominant expression axis, and ancestry remains the larger block after separation from ClockPC1. (a) Pairwise Spearman correlations among the 19 home-site bioclimatic variables. (b) Five-fold cross-validated R² for environmental principal components alone, genomicPC1–10 alone, ClockPC1 alone, the 13-variable structural set (genomicPC1–10, latitude, longitude and ClockPC1), and the structural and environmental sets combined. (c) Variance partitioning of ExprPC1 under the three-block allocation with geography assigned to the environmental block: 5.0% unique to genomicPC1–10, 19.8% to ClockPC1, 2.6% to environment and geography, 7.4% shared and 65.2% unexplained. The primary two-block model combines genomicPC1–10 and ClockPC1 into a structural block that uniquely explains 26.0% (Table S3).

### Circadian and developmental covariates

Because collection timing inevitably varies across an experiment of this size, analyses included ClockPC1, the first principal component of nine core circadian clock genes (CCA1, LHY, TOC1/PRR1, PRR7, PRR9, GI, ELF3, ELF4 and LUX; McClung 2006). Its negligible correlations with climate (maximum |Spearman ρ| = 0.063, all P > 0.1) support its use as a circadian rather than an environmental covariate (Fig. S2). ClockPC1 could in principle also absorb genotypic differences in clock phase, but it was not detectably heritable (h² = 0.02, interval 0.00–0.13, P = 0.15; Table S11), which bounds a heritable clock contribution without excluding one; we therefore carry it as a circadian covariate. A broader alternative, CircPC1, was derived identically from the 347 feature genes selected by ChronoGauge’s sequential feature selection (Reynolds et al. 2025), only two of which are shared with ClockPC1. CircPC1 uses that gene set alone and is therefore a covariate axis rather than an estimate of sampling time. As an external check on the covariate, we also ran the ChronoGauge predictor itself on these libraries, using 84 of its 100 released sub-predictors and aggregating their estimates by circular mean (Methods S1; Fig. S7). StagePC1 summarized 20 available floral-transition markers from a 21-gene set, with its sign aligned so higher values indicate greater progression toward bolting.

### Co-expression clustering

We hierarchically clustered standardized expression for 24,175 genes across 664 accessions using Pearson distance (1 − *r*), average linkage and a dendrogram cut height of 0.98. Clusters with <30 genes were unassigned, yielding 23 modules and 335 unassigned genes. Module eigengenes are first principal components of standardized member expression. Functional labels were assigned from enriched GO:BP terms matching proliferation, defense or photosynthesis keywords. Robustness tests varied the cut height from 0.93 to 0.99, removed low-membership genes, added high-membership non-members and forced two to six submodules (Methods S1).

### Functional enrichment, promoter motifs and hub genes

Gene Ontology enrichment used g:Profiler (Kolberg *et al*. 2023) with a Benjamini–Hochberg threshold of q < 0.05; broad non-specific terms were retained in the full enrichment and the terms displayed in Fig. 2(b) are a selection of the more specific ones. Module genes were partitioned into cell cycle, ribosome and other categories by GO membership, and four transcription factor motifs were queried in the 2 kb upstream of each transcription start site, with enrichment between categories tested by Fisher’s exact test. Hub genes were the top 1% of members by module membership score. Motif sequences, GO term lists and the transcription factor family assignment are given in Methods S1.

**Fig. 2.**
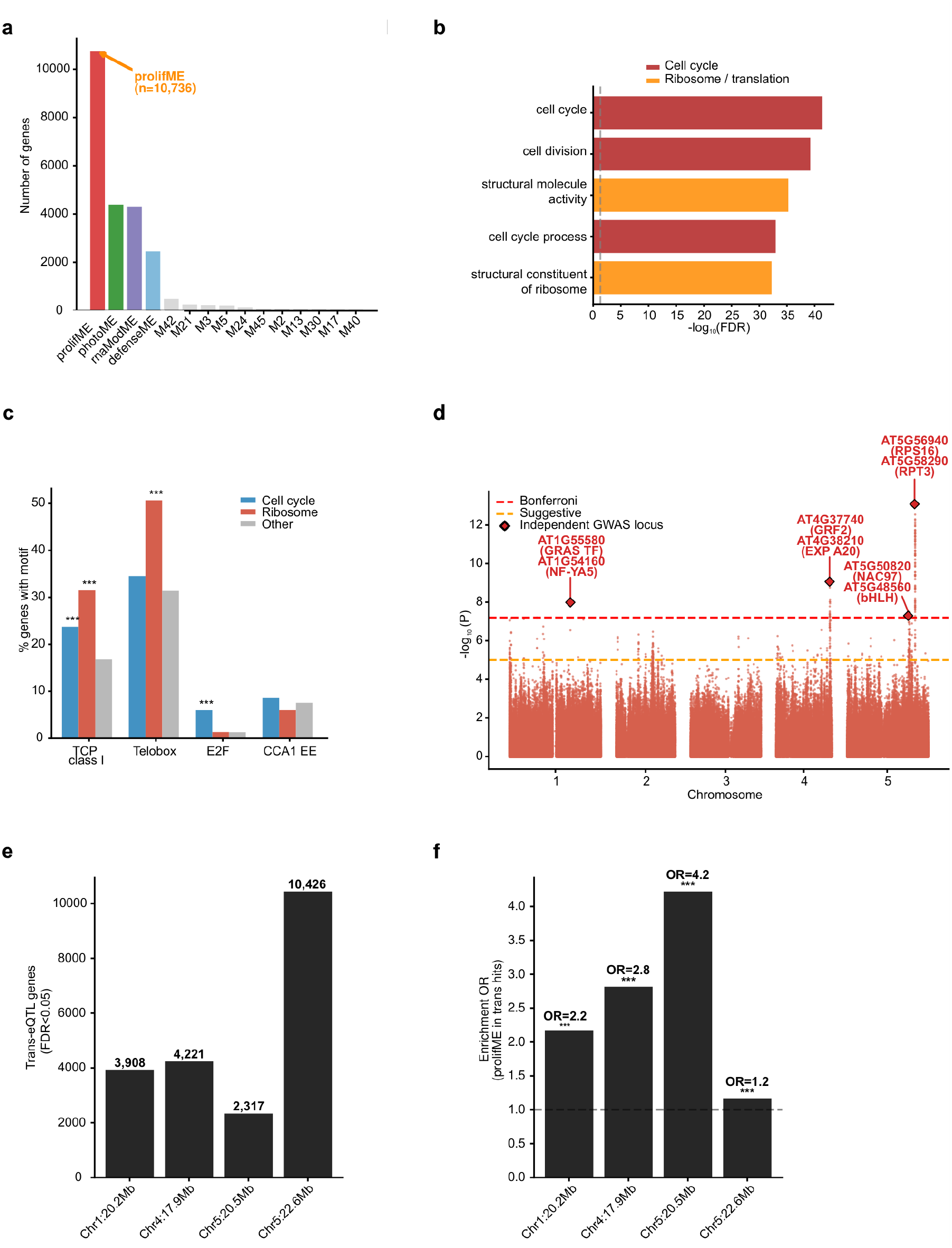
The dominant expression axis is associated with a proliferation module with a bipartite motif structure and four loci showing widespread trans associations. (a) Sizes of the co-expression modules, including prolifME (10,736 genes). (b) Selected GO biological-process enrichments for prolifME. (c) Frequencies of TCP class I, telo-box, E2F and Evening Element motifs in the 2 kb upstream of the transcription start site for cell-cycle, ribosome and other prolifME genes. Asterisks denote Fisher’s exact tests against the other-gene category. (d) Genome-wide association for the prolifME eigengene, with suggestive and Bonferroni thresholds indicated. (e) Number of significant trans-associated genes per lead SNP (FDR < 0.05). (f) Enrichment of prolifME genes among those trans-associated genes.

### Variance partitioning

Unique and shared contributions of environmental and structural blocks followed Peres-Neto et al. (2006). Five-fold cross-validated ridge-regression R² provided the predictive comparison in Fig. 1(b); its 13-variable structural set comprised genomicPC1–10, latitude, longitude and ClockPC1. Single-axis partitions used ExprPC1 (8.0% of total expression variance), so their percentages refer to that axis. We varied the assignment of latitude, longitude and ClockPC1 among predictor blocks (Table S3). Transcriptome-wide partitioning used adjusted-R² redundancy analysis of all 24,175 genes and an expression-permutation null. The primary 531-accession model contained genomicPC1–10 and ClockPC1 in the structural block and 19 bioclimatic variables, eight soil variables, latitude and longitude in the environmental block. Soil-excluded sensitivity analyses used the same 531 accessions and the 663 accessions with complete climate and coordinate records (Table S12). Further controls included 200 response permutations within each of five geographic subsets, leave-one-admixture-group-out analyses, explicit admixture-group labels, and CircPC1 and StagePC1 as additional blocks (Tables S3–S8).

### SNP heritability

SNP heritability of each module eigengene and of the leading expression axes was estimated by restricted maximum likelihood on a genomic relationship matrix built from the 1001 Genomes genotypes. Variance components were obtained by profiling the restricted likelihood over the eigendecomposition of that matrix, and intervals are 95% profile intervals. Significance was assessed by referring the likelihood-ratio statistic to a null built from 200 permutations of the phenotype, because its asymptotic boundary distribution is unreliable for these traits (Methods S1). Conditional estimates include genomicPC1–10 as fixed effects (Table S11).

### Genome-wide association and *trans*-eQTL analysis

Association testing used the prolifME eigengene as a quantitative trait in a mixed linear model with a genomic relationship matrix and genomicPC1–10 as fixed-effect covariates (GCTA; Yang *et al*. 2011). Independent signals were defined by a Bonferroni threshold and LD-based clumping in plink2 (r² < 0.1 within 1 Mb), and candidate genes were taken within ±500 kb of each lead SNP from the TAIR10 annotation. For each lead SNP, *trans* effects were tested by regressing every gene’s z-scored expression on genotype with genomicPC1–10 as covariates, restricted to gene–SNP pairs on different chromosomes or more than 1 Mb apart, with Benjamini–Hochberg correction applied per locus. Enrichment of module genes among *trans* targets was assessed by Fisher’s exact test. Allele frequencies at lead SNPs were computed per admixture group and ordered by mean latitude.

### Phenotype association and benchmarking

For the four growth-related phenotypes, partial Pearson correlations were computed after regressing each trait and eigengene on genomicPC1–10, latitude, longitude and ClockPC1; StagePC1 and CircPC1 were added in sensitivity tests. The flowering-time estimate in Fig. S5(b) uses the primary covariate set of genomicPC1–10, latitude, longitude and ClockPC1; Table S6 reports the sensitivity analysis with StagePC1 added. Control sets tested module composition: cell-cycle and ribosomal subsets within prolifME and the least variable genes inside and outside it, each compared with matched draws from the appropriate gene pool (Methods S1). Matched-set nulls used up to 1,000 draws, and phenotype-label tests used 1,000 permutations. Latitudinal directional coherence was the fraction of module genes with negative coefficients after adjustment for longitude and ClockPC1, compared with size- and expression-matched external sets.

### Cross-species projection

Orthogroups among *Arabidopsis thaliana*, *Oryza sativa* and *Zea mays* were inferred with OrthoFinder v2.5 from Ensembl Plants and Phytozome proteomes (Emms & Kelly 2019). *Arabidopsis* gene contributions were correlations with prolifME across 665 accessions. Rice scores were derived from a compiled matrix of 8,456 public RNA-seq runs; the processed matrix and complete run-accession manifest will accompany the analysis code in the Zenodo archive. Maize scores were derived from 1,913 samples (Kremling et al. 2018). For both species, scores were estimated independently as the first principal component of orthologous genes within that species, and contributions were correlations with that score. Score signs were fixed within species to give positive mean contributions. Alignment was computed between independently estimated contribution vectors. Significance used 10,000 orthogroup-label permutations and a sign-randomization control; specificity was tested within orthogroups assigned to each module.

### Statistical analysis

Analyses were performed in Python 3.10 with NumPy, SciPy, pandas, scikit-learn and statsmodels. All *P* values are two-sided unless stated otherwise, and multiple testing was controlled by the Benjamini–Hochberg procedure where indicated. Full protocols and all sensitivity analyses are given in Methods S1.

## Results

### Ancestry predicts the dominant expression axis more strongly than home-site environment

We first compared ancestry and home-site environment as predictor blocks on the leading expression axes. In five-fold cross-validation on ExprPC1, environmental principal components alone reached a CV-R² of 0.013 and genomic principal components alone 0.079. ClockPC1 alone reached 0.197, and the 13-variable structural set (genomicPC1–10, latitude, longitude and ClockPC1) reached 0.262; adding the environmental block increased CV-R² only to 0.264 (Fig. 1b). Thus, most of the predictive advantage of the full structural set came from the circadian covariate rather than ancestry. Cross-validation ranks predictor sets but does not separate their contributions, which we addressed by variance partitioning.

We partitioned the dominant expression axis (ExprPC1), which accounts for 8.0% of total expression variance (Table S4). In the primary two-block model, the structural block uniquely explained 26.0% of ExprPC1 and environment 2.6%, with 6.1% shared and 65.2% unexplained (Table S3). A three-block model that separated ancestry, ClockPC1 and environment assigned 5.0% uniquely to ancestry, 19.8% to ClockPC1 and 2.6% to environment, with 7.4% shared (Fig. 1c). Four assignments of geography and the circadian covariate preserved the ancestry–environment ordering (Table S3). In the genomic-only specification, ancestry uniquely explained 8.8% of ExprPC1, compared with 1.4% for environment. The structural and environmental blocks shared 6.1% because both covary with geography, and genomicPC1–10 strongly predicted latitude and longitude (Methods S1). The same structural–environment ordering held across all ten leading expression axes (Table S4; Fig. S1b). The ordering is consistent across specifications; its magnitude is not.

The ordering also held within regions, where climate varies over a narrower range. We repeated the two-block partition in five geographic subsets containing 53–126 accessions. Because each regional model had 23 predictors, we expressed each unique fraction relative to its own null from 200 response permutations (Fig. S1c). The structural block exceeded its null in all five regions, from 1.04-fold in Sweden to 3.91-fold in Central Europe (median 2.11-fold), and remained significant after Benjamini–Hochberg correction in Germany, Spain, Western Europe and Central Europe (q = 0.025). The environmental block had a median ratio of 0.58 and was significant in no region (q ≥ 0.129). Sweden was the only region in which the environmental ratio exceeded the structural ratio (1.70 versus 1.04), but neither block differed from its null there. Restricting geographic range weakened both signals without reversing their ordering where either block was distinguishable from its null.

The ancestry–environment difference was smaller across the complete expression matrix. Redundancy analysis of all 24,175 genes in the 531 accessions with complete climate and soil records assigned 6.1% of expression variance uniquely to the full structural block and 2.2% to environment, with 3.6% shared and 88.1% unexplained (Table S12). Restricting the structural block to genomicPC1–10 left 2.6% for ancestry and 2.0% for environment. Excluding the eight soil variables reduced the environmental fraction to 1.5% in the same accessions and to 1.43% in the 663 accessions with complete climate and coordinate records; the corresponding ancestry fractions were 2.9% and 2.87%. The ancestry advantage was concentrated on the leading expression axes and smaller across the transcriptome as a whole.

Discrete subpopulations also did not account for the signal. GenomicPC1–10 retained explanatory power after adjustment for admixture-group identity, whereas group identity added little after adjustment for the genomic components. Removing each group changed the structural fraction only modestly, and the ordering held within seven of eight groups with at least 40 accessions (Table S8; Methods S1). Circadian and developmental proxies likewise changed the estimated fractions but not their ordering. CircPC1, derived from 347 ChronoGauge genes, shared only two genes with ClockPC1 but was strongly correlated with it and added explanatory power for ExprPC1. With ClockPC1, CircPC1 and StagePC1 all conditioned on at once, the incremental variance explained by ancestry was 4.9 times that explained by climate and soil for ExprPC1, and 7.1 times for prolifME (Table S7). A four-block partition that instead divides total variance among ancestry, ClockPC1, StagePC1 and environment gives smaller fractions. For ExprPC1, ClockPC1 took 19.1%, StagePC1 6.5%, ancestry 6.0% and climate and soil 3.1% (Table S6), so the circadian covariate is the largest single block and ancestry ranks third, behind StagePC1; StagePC1 had to be derived from expressed floral-transition markers because no independent staging record was available. For prolifME the order is reversed, with ancestry at 5.2%, ClockPC1 at 4.4%, environment at 3.9% and StagePC1 at 1.9% (Table S6). The incremental and partitioned estimates are not comparable in size, but ancestry exceeds environment in both.

### A heritable proliferation program underlies the axis and is associated with four loci

We next asked which co-expression module carried ExprPC1. One broad module eigengene correlated with ExprPC1 at ρ = 0.858, and ExprPC1 and ExprPC3 together explained 92.7% of its variation (Fig. S3a,b). The module contains 10,736 genes, approximately 44% of the expressed transcriptome. Cell-cycle regulation and ribosome biogenesis were its clearest specific enrichments, whereas stress and environmental-sensing terms were not enriched (Fig. 2a,b; Fig. S3g). We call this eigengene prolifME and use defenseME, photoME and rnaModME as comparators (Fig. 2a; Fig. S3c–e). Removing ribosomal genes left prolifME almost unchanged, indicating that ribosomal abundance alone does not define the module (Fig. S3f).

Promoter motifs separated the cell-cycle and ribosome components of prolifME. E2F motifs were enriched among cell-cycle genes, whereas TCP class I and telo-box motifs were enriched among ribosome genes (Fig. 2c), a contrast that held across all four modules (Fig. S3h). These factor families were absent from the top prolifME hub genes (Fig. S3i). The motif structure is consistent with known control of cell division and protein synthesis and nominates these factors as upstream regulators rather than central module hubs (Vandepoele et al. 2002; Trémousaygue et al. 2003; Sozzani et al. 2010). Because telo-box-associated ribosomal genes can be circadian-regulated, the primary models included ClockPC1 to account for circadian expression variation; replacing ClockPC1 with the independently defined CircPC1 retained the ancestry–environment ordering (Table S7).

The prolifME eigengene was stable to changes in module definition. Re-running the clustering recovered all 23 modules and their eigengenes. Varying the cut height changed membership substantially, but every resulting eigengene remained strongly correlated with the original (|r| ≥ 0.956; Table S9). Removing half the member genes, or sampling a random 10% of them, reproduced the eigengene at |r| ≥ 0.995. Eigengene-level results were insensitive to the module boundary, although exact gene membership was not (Methods S1).

ProlifME was also heritable. Its SNP-heritability estimate was h² = 0.19 (95% profile interval 0.12–0.29; Table S11) and fell only to 0.17 after adjustment for genomicPC1–10. None of 200 phenotype permutations reached the observed likelihood-ratio statistic (empirical P = 1/201 = 0.005), although simulations showed that the point estimate remains imprecise at this sample size (Methods S1). The other modules had overlapping intervals: h² = 0.26 for defenseME, 0.21 for photoME and 0.30 for rnaModME. ClockPC1 was not detectably heritable (h² = 0.02, interval 0.00–0.13, P = 0.15), which bounds a heritable clock contribution without excluding one. ChronoGauge circadian-time estimates for these libraries fell in a narrow window (median 6.9 h, interquartile range 5.5–8.2 h; Fig. S7a) and tracked ClockPC1 closely (Spearman ρ = 0.866, P < 1 × 10⁻²⁰⁰, n = 727; Fig. S7b). Because ChronoGauge is itself estimated from expression, that agreement shows that ClockPC1 tracks circadian state but cannot distinguish collection-time variation from accession-specific differences in circadian phase or period.

Genome-wide association testing identified four loci associated with prolifME after adjustment for genomicPC1–10, each separating the eigengene by genotype (Fig. 2d; Fig. S4a–d; Table S2). The loci were near NF-YA5, GRF2, NAC97, and a region containing RPS16 and RPT3 (Kim et al. 2003; Li et al. 2008; Table S1). Each locus was associated in trans with 2,317–10,426 genes at FDR < 0.05, and these genes were enriched in prolifME (odds ratio 1.2–4.2; Fig. 2e,f). Because the loci were selected through the module eigengene, this enrichment is partly expected by construction. Seven of the eight nominated genes belong to prolifME, including at least one at each locus; RPS16 at the Chr5:22.6 Mb locus belongs to rnaModME (Table S1).

Selection-aware estimates showed that the four loci had modest predictive value. The within-sample adjusted R² was 15.45%, but phenotype permutation attributed 11.49 percentage points to selection alone, leaving an excess of 3.96 points (Table S5). Repeating SNP selection within each cross-validation fold retained 2.68% out of sample. Allele frequencies were strongly population-structured; at Chr5:22.6 Mb, the high-prolifME A allele was least frequent in northern Sweden (Fig. S4e–h). The four loci therefore contribute to prolifME but do not account for most of its variation.

The module’s members are spread across the genome rather than concentrated in one region. ProlifME genes were more dispersed than genes in size-matched random sets, with a median nearest-neighbor distance of 4,090 bp compared with 3,870 ± 26 bp in matched sets. We residualized expression on genomicPC1–10 together with latitude and longitude, which reduced the variance explained by the module’s first principal component only from 10.1% to 9.4%. The axis survives removal of the leading ancestry coordinates and geography, and its members are not confined to one haplotype block.

### The proliferation program is latitudinally biased and predicts growth phenotypes

An expression axis matters biologically only if it reaches the whole-plant phenotype, so we tested the module against geography and archived phenotypes. ProlifME showed the strongest latitudinal bias among the four modules. Expression decreased with latitude for 72.0% of prolifME genes and 62.7% for defenseME, whereas rnaModME was biased in the opposite direction (35.4% declining) and photoME showed no significant bias (Fig. 3a; Fig. S5a). The prolifME fraction exceeded matched gene-set nulls, whereas its adjusted mean coefficient was not significant (P = 0.083). Higher prolifME was also associated with lower defenseME (r = −0.261, n = 664; Fig. 3b). Flowering was earlier in the same direction but did not reach significance under the declared covariates (r = −0.076, P = 0.054, n = 648; Fig. S5b). It reached r = −0.131 (P = 8.3 × 10⁻⁴) only when the developmental-stage covariate was added (Table S6). Because StagePC1 is itself a transcriptional readout of the floral transition (r = −0.378 with flowering time), we do not count that association as independent evidence. The pattern is consistent with a transcriptome-level growth–defense contrast (Huot et al. 2014; Züst & Agrawal 2017).

**Fig. 3.**
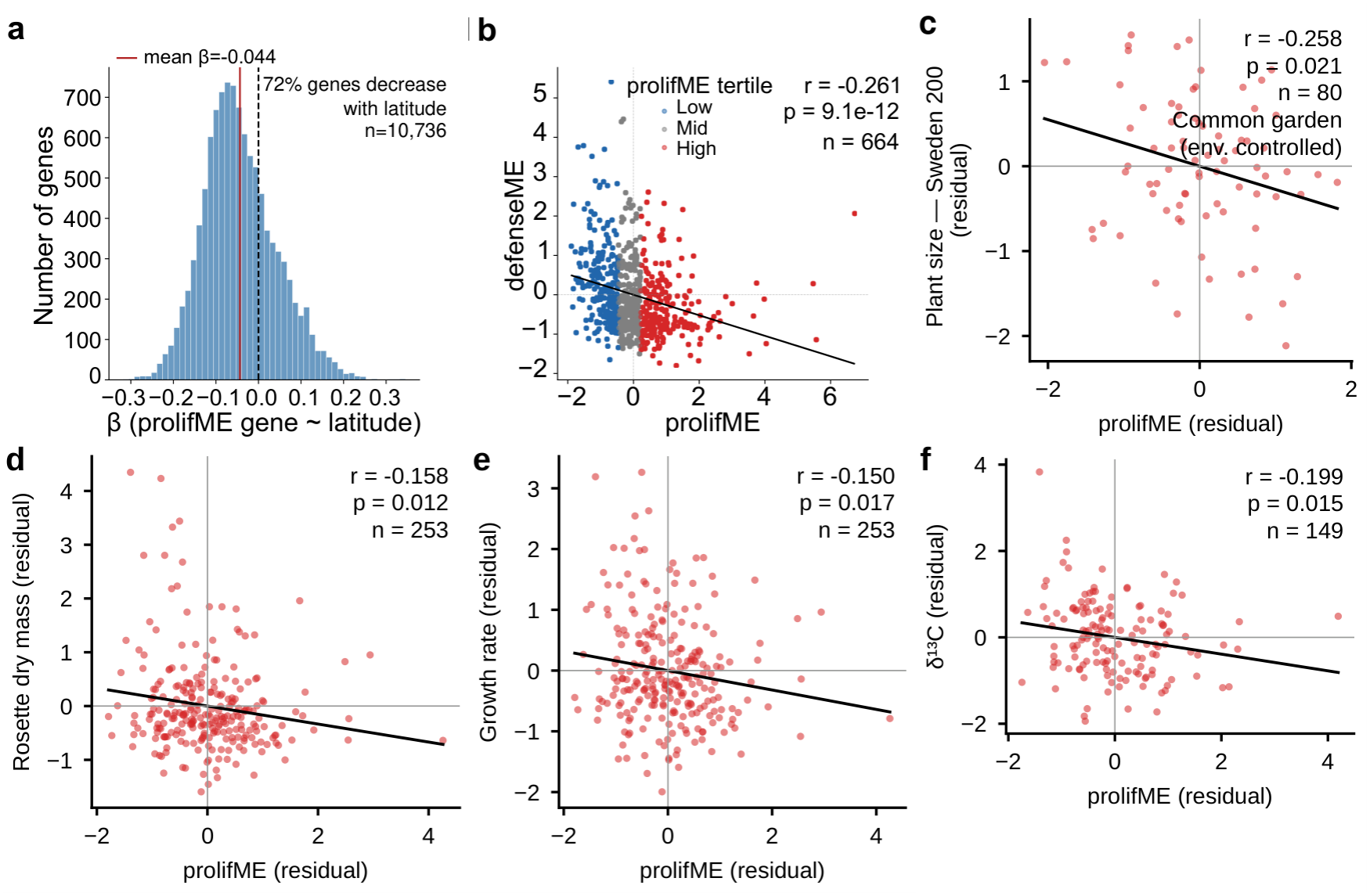
prolifME is geographically structured and is associated with growth-related phenotypes. (a) Distribution of gene-level latitude coefficients within prolifME after adjustment for longitude and ClockPC1; 72.0% are negative and the mean coefficient is −0.044. Because module genes are not independent, inference uses size- and expression-matched gene sets drawn from outside the module rather than a binomial test (Fig. S5a; Methods S1). (b) Relationship between prolifME and defenseME across accessions (n = 664). (c) Swedish common-garden plant size against prolifME (n = 80). (d, e) Rosette dry mass and growth rate against prolifME under controlled conditions (n = 253). (f) Leaf carbon isotope composition (δ¹³C) against prolifME (n = 149). Panels (c–f) show residuals after adjustment for genomicPC1–10, latitude, longitude and ClockPC1.

The phenotypes were identified through an initial screen, and we quantified that selection explicitly. Of 535 AraPheno traits, 461 had sufficient overlap with the expression data. Rosette dry mass and flowering time at 10 °C survived screen-wide Benjamini–Hochberg correction, and growth rate, common-garden size and leaf carbon isotope composition (δ¹³C) also ranked among the eight strongest associations (Table S13). The screen documents how the traits were selected, whereas the adjusted models provide the relevant effect-size estimates.

Higher prolifME was associated with smaller plants in two independent experiments and with slower growth in the controlled experiment. In a Swedish common garden, higher prolifME was associated with smaller plants (r = −0.258, P = 0.021, n = 80; Ågren & Schemske 2012; Fig. 3c). Under controlled conditions, higher prolifME was associated with lower rosette dry mass (r = −0.158, P = 0.012) and slower growth (r = −0.150, P = 0.017; n = 253 for both; Vasseur et al. 2018; Fig. 3d,e). Higher prolifME was also associated with lower leaf δ¹³C (r = −0.199, P = 0.015, n = 149; Fig. 3f). Lower δ¹³C corresponds to greater carbon isotope discrimination and therefore to lower intrinsic water-use efficiency. Although the individual effects were modest, all four correlations were negative. Phenotype-label permutations supported all four associations (P = 0.016–0.039), and adjustment for StagePC1 retained all four and strengthened the common-garden association to r = −0.316 (P = 0.0043; Table S6). Because the ancestry components are removed in these models, the associations are independent of the ancestry result.

Composition controls localized the phenotype signal within the module. Cell-cycle and ribosomal subsets tracked prolifME across the three traits (r = −0.14 to −0.21), whereas the least variable genes inside it were weaker (r = +0.04 to −0.08; Fig. S5c). The two functional subsets were not distinguishable from random prolifME draws, and a low-variance set outside prolifME reproduced the common-garden association. The phenotype signal follows expression variability within this broad module rather than the cell-cycle–ribosome split itself.

### *Arabidopsis* gene weights correlate with independent rice and maize profiles

We then asked whether the gene-weight profile of the *Arabidopsis* axis correlates with independently estimated profiles in other species. Across all shared orthogroups, *Arabidopsis* gene contributions correlated with those in rice (r = 0.446) and maize (r = 0.379; Fig. 4a). Neither sign randomization nor 10,000 orthogroup-label permutations reproduced correlations of this magnitude (P < 1 × 10⁻⁴; Fig. 4c), and alignment was stronger among strict one-to-one orthologs (r = 0.538 in rice and 0.461 in maize; Fig. S6).

**Fig. 4.**
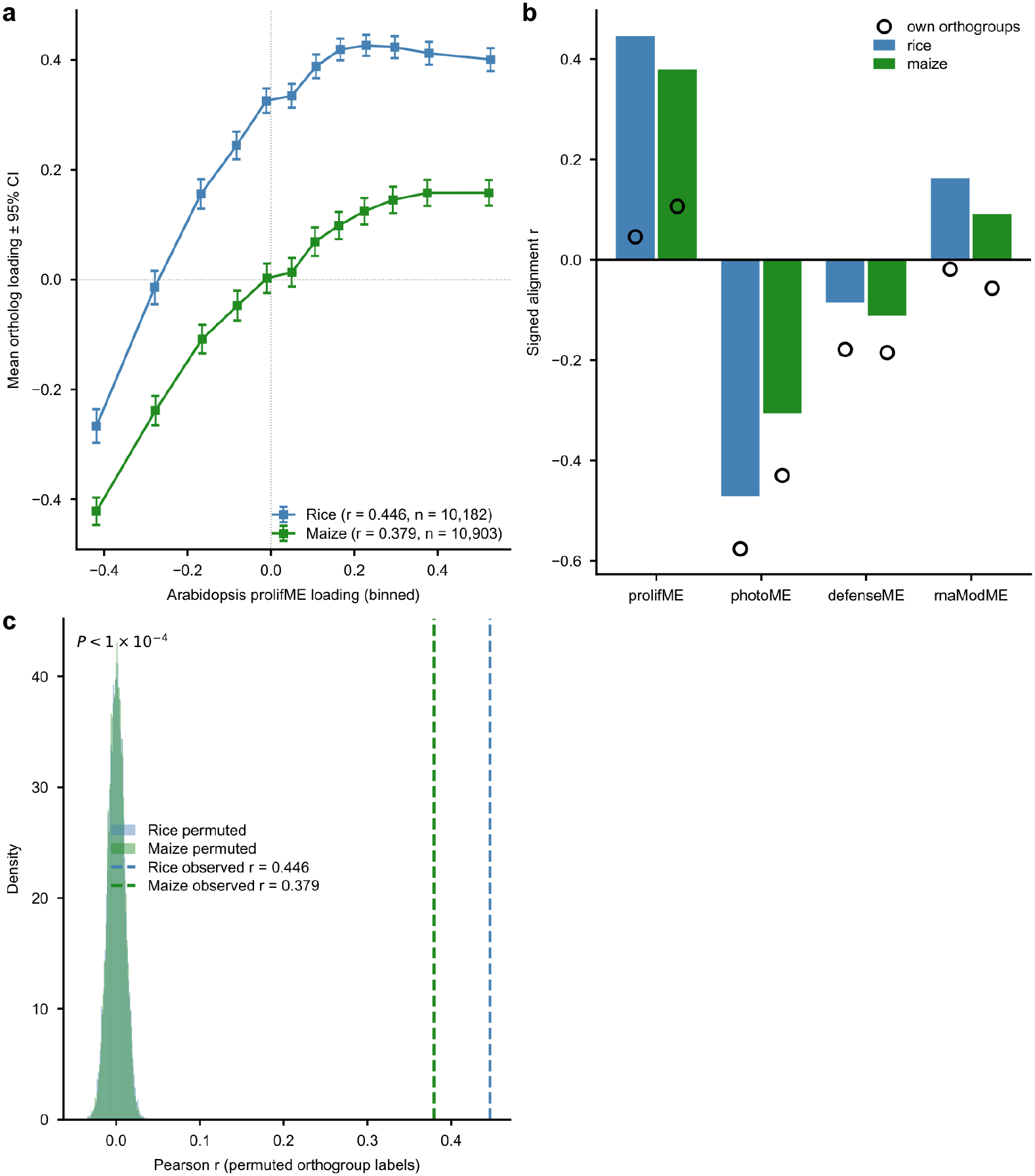
Arabidopsis gene weights correlate with independent rice and maize profiles. (a) Arabidopsis prolifME gene contributions, binned along the horizontal axis, against mean orthologous contributions estimated independently in rice and maize; points show bin means and 95% confidence intervals. (b) Signed alignment for all four modules. Filled bars use all shared orthogroups, and open circles restrict each module to its own orthogroups. The photosynthetic alignment is retained under this restriction, whereas the proliferative alignment approaches zero. (c) Null distributions from 10,000 orthogroup-label permutations, with the observed correlations indicated. The minimum attainable empirical P value is 1/10,001 (< 1 × 10⁻⁴).

Module-level analyses localized the cross-species signal. PhotoME aligned with the opposite sign in both species (r = −0.471 in rice and −0.306 in maize), whereas defenseME and rnaModME aligned only weakly (|r| ≤ 0.163; Fig. 4b). Restricting the calculation to each module’s own orthogroups preserved the photosynthetic alignment (r = −0.576 in rice and −0.430 in maize) but nearly abolished the proliferative alignment (r = 0.046 and 0.107; Table S10). A paired bootstrap identified prolifME as the top-aligned module in every maize resample but in no rice resample. The cross-species correspondence is carried largely by the photosynthetic pole and does not imply conserved gene ordering within prolifME.

## Discussion

Studies of expression variation in natural plant populations have usually focused on environmental responses of individual genes (Des Marais & Juenger 2010; Lasky et al. 2012). We asked a different question: what predicts the major axes of variation among accessions grown under one regime? Genomic ancestry explained more of the dominant axis than home-site climate and soil in every block specification, and that axis carries a heritable, cell-cycle- and ribosome-enriched module that tracks growth phenotypes. Because the accessions were grown under a single condition, the result concerns differences that persist in a common environment rather than acute plasticity.

The ancestry–environment ordering was stable across sensitivity analyses. It survived alternative assignments of geography and circadian covariates, removal of individual admixture groups, adjustment for explicit group labels and restriction to single geographic regions, where it held in four of the five regions and in seven of the eight admixture groups. If anything the within-region comparison understates ancestry, because genomicPC1–10 summarizes continental structure and may miss fine-scale relatedness inside a region. Ancestry and environment nevertheless share substantial geographic signal, and a climate-only analysis can assign that shared variance to climate unless population structure is modeled explicitly (Peres-Neto et al. 2006). The claim is also scale-dependent: ancestry is the stronger predictor of the leading axes, whereas across all 24,175 genes neither block dominates (2.6% versus 2.0%; Table S12). Local adaptation at individual loci and traits is well established in *Arabidopsis* (Hancock et al. 2011; Fournier-Level et al. 2011; Lasky et al. 2012), and particular genes on an ancestry-associated axis may remain climate-responsive. This analysis separates the organization of a major expression axis from gene-specific environmental response; it does not establish adaptation.

The biological content of ExprPC1 is a proliferation-associated state in which cell-cycle and ribosome genes vary together, a contrast recurring across eukaryotic transcriptomes (Gasch et al. 2000; Oldham et al. 2008; Zhang et al. 2022), and its bipartite promoter-motif structure connects this population-level axis to established plant control of cell division and protein synthesis. Because independent halves of the gene set reconstruct the axis at *r* = 0.998, the 65.2% residual reflects variation the predictor blocks do not capture rather than noise in the response. The heritability of prolifME is not a restatement of the ancestry association: adjustment for genomicPC1–10 changed the estimate little. ExprPC1 was heritable as well (*h*² = 0.21, 95% profile interval 0.12–0.32; Table S11), so prolifME is distinctive for dominating ExprPC1 and tracking growth phenotypes, not for being the most heritable feature. The four loci sit alongside earlier expression-QTL work. Trans-acting loci that affect hundreds to thousands of transcripts at once are well documented in biparental *Arabidopsis* populations (West et al. 2007; Lensink et al. 2025), whereas in panels of natural accessions per-gene mapping has recovered mostly cis effects, with trans signals a minority and individually weak (Zan et al. 2016). Using a module eigengene as the mapped trait collects many small, correlated trans effects into one test, and so reveals associations resembling eQTL hotspots in an accession panel. The same feature limits interpretation. Association in accessions is confounded by population structure, which is the subject of this paper rather than a nuisance set aside, and the loci were selected through the eigengene itself, which is why we report selection-aware and out-of-sample estimates, with 2.68% retained out of sample. Hotspot positions from biparental crosses and from accession panels are not directly comparable, and we make no claim of correspondence; the loci are markers rather than causal variants, distributed cis-eQTLs remain a possible source of the ancestry association, and the assignment of *RPS16* to rnaModME hints that the Chr5:22.6 Mb locus links the proliferation and RNA-modification programs.

The phenotype results give prolifME an independent biological interpretation. Higher prolifME was associated with smaller size and slower growth in two experiments and with lower leaf δ¹³C, and all four associations exceeded phenotype-label nulls. Because the models remove the ancestry components, they are independent of the ancestry result rather than extensions of it. The growth–defense trade-off is usually studied as an induced response, in which a plant is challenged and growth slows. These data describe something different: a standing difference among accessions grown together with no challenge applied. ProlifME and defenseME vary in opposite directions across the panel (*r* = −0.261, *n* = 664), each is heritable (*h*² = 0.19 and 0.26), and both decline with latitude in the same direction (72.0% and 62.7% of their genes). Together these results raise the possibility that accessions differ in a partly heritable transcriptional balance between growth- and defense-associated programs, and such standing variation may be relevant to breeding and synthetic biology. Two limits bound that reading. DefenseME is a transcriptional module rather than measured resistance, and no pathogen assay or fitness measurement is reported here; and the two eigengenes are not independent coordinates of a single axis, so their negative correlation does not establish a causal trade-off.

Overall, the gene weights defining the *Arabidopsis* axis were correlated with independently derived profiles in rice and maize, with the correspondence carried largely by the photosynthetic pole and gene ordering within prolifME not conserved. The defense pole did not align in either species (|*r*| ≤ 0.163; Fig. 4b), so the growth–defense contrast seen within *Arabidopsis* is not recovered in these crops; whether that reflects a different organization of the trade-off, their domestication history, or the narrower sampling of the compiled expression panels, these data cannot say. The discovery data come from one tissue, developmental stage and growth condition and were aligned to the Col-0 reference, yet the ancestry ordering persisted after developmental and circadian adjustment, and removing the genetically most divergent relict accessions increased rather than reduced the ancestry fraction (Table S8). Genomic ancestry therefore predicts a heritable proliferation-associated expression axis more strongly than home-site climate and soil, identifying a phenotypically anchored level of transcriptome organization that complements gene-by-gene models of environmental response.

## Acknowledgements

We thank members of the Lee and Busch laboratories for critical discussions. This research was supported by the National Research Foundation of Korea (NRF), funded by the Korean government (MSIT, MOE) (RS-2025-21882992, RS-2018-NR031069 and RS-2026-25470625), and by the Yonsei University Research Fund (2025-22-0153, 2026-22-0199) to S.L.

## Author contributions

Sanghwa Lee (here after S.L.) conceived the study and designed the experiments. Soomin Lee, C.L., and D.G. analyzed the data. S.L. supervised work and provided funds and resources. S.L. wrote the manuscript with input of all the authors. S.R., H.J., W.B., and T.P.M. commented the paper.

## Data availability

Primary data are public: *Arabidopsis* transcriptomes (GEO GSE80744), 1001 Genomes genotypes, AraPheno phenotypes, WorldClim v2.1 climate, SoilGrids soils and maize expression data (Kremling et al. 2018).

## Supporting Information

Methods S1 Detailed protocols and sensitivity analyses.

**Fig. S1.**
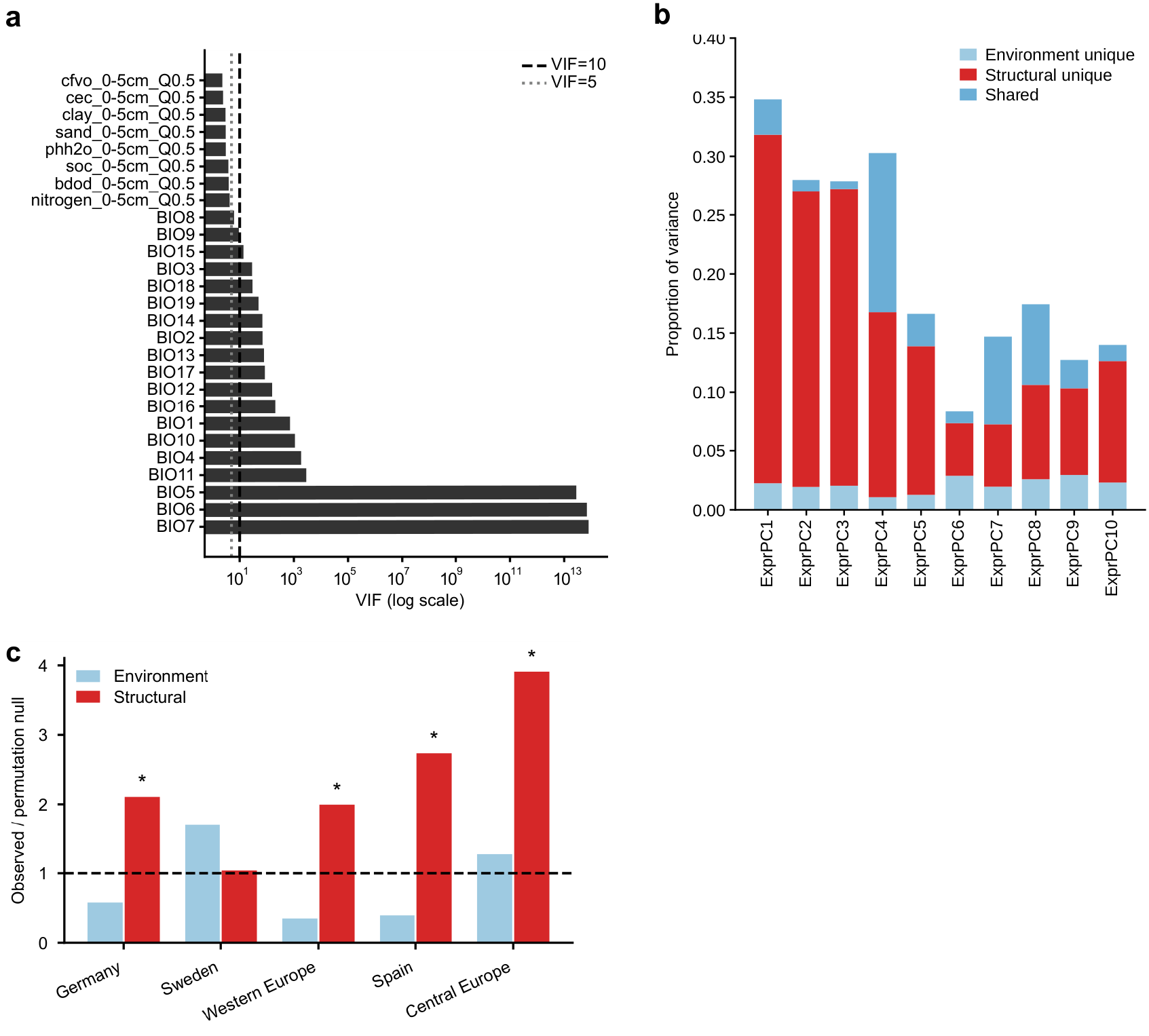
Environmental predictor collinearity and the scale dependence of the ancestry–environment comparison. (a) Variance inflation factors for the 19 bioclimatic and eight soil variables, on a log scale; the dashed and dotted lines mark VIF = 10 and VIF = 5. Most bioclimatic variables exceed a VIF of 10, several by many orders of magnitude, whereas none of the soil variables do. (b) Variance partitioning of each of the ten leading expression principal components (ExprPC1–10) into the unique contribution of the environmental block (the 19 bioclimatic and eight soil variables), the unique contribution of the structural block (genomicPC1–10, latitude, longitude and ClockPC1), and the fraction they share, showing that the ordering in Fig. 1c is not specific to ExprPC1. (c) Variance partitioning within five geographic regions (Sweden n = 64, Spain n = 101, Germany n = 53, Central Europe n = 126, Western Europe n = 77), expressed as the observed unique fraction divided by the mean of a region-specific permutation null (200 permutations of the expression response with that region’s observed predictor matrix held fixed). The dashed line marks the null expectation and asterisks mark P ≤ 0.05 after Benjamini–Hochberg correction across the five regions. Raw unique fractions are not comparable between regions because each subset holds 53 to 126 accessions against 23 predictors.

**Fig. S2.**
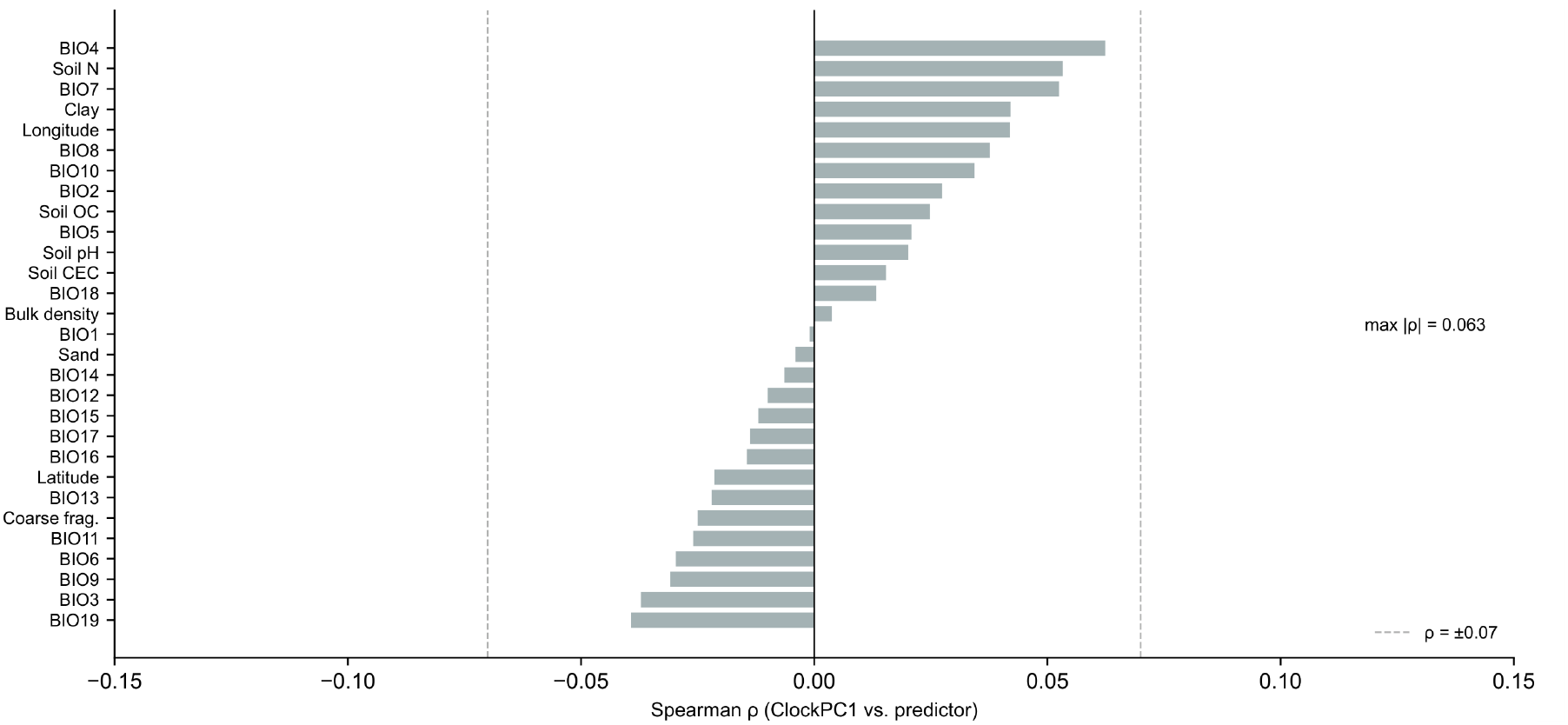
The circadian covariate is unrelated to home-site climate. ClockPC1, the first principal component of nine clock-associated genes (CCA1, LHY, TOC1/PRR1, PRR7, PRR9, GI, ELF3, ELF4 and LUX), plotted against each bioclimatic, soil and geographic predictor. The largest Spearman correlation is |ρ| = 0.063, with all P > 0.1, which supports treating ClockPC1 as a circadian rather than an environmental covariate.

**Fig. S3.**
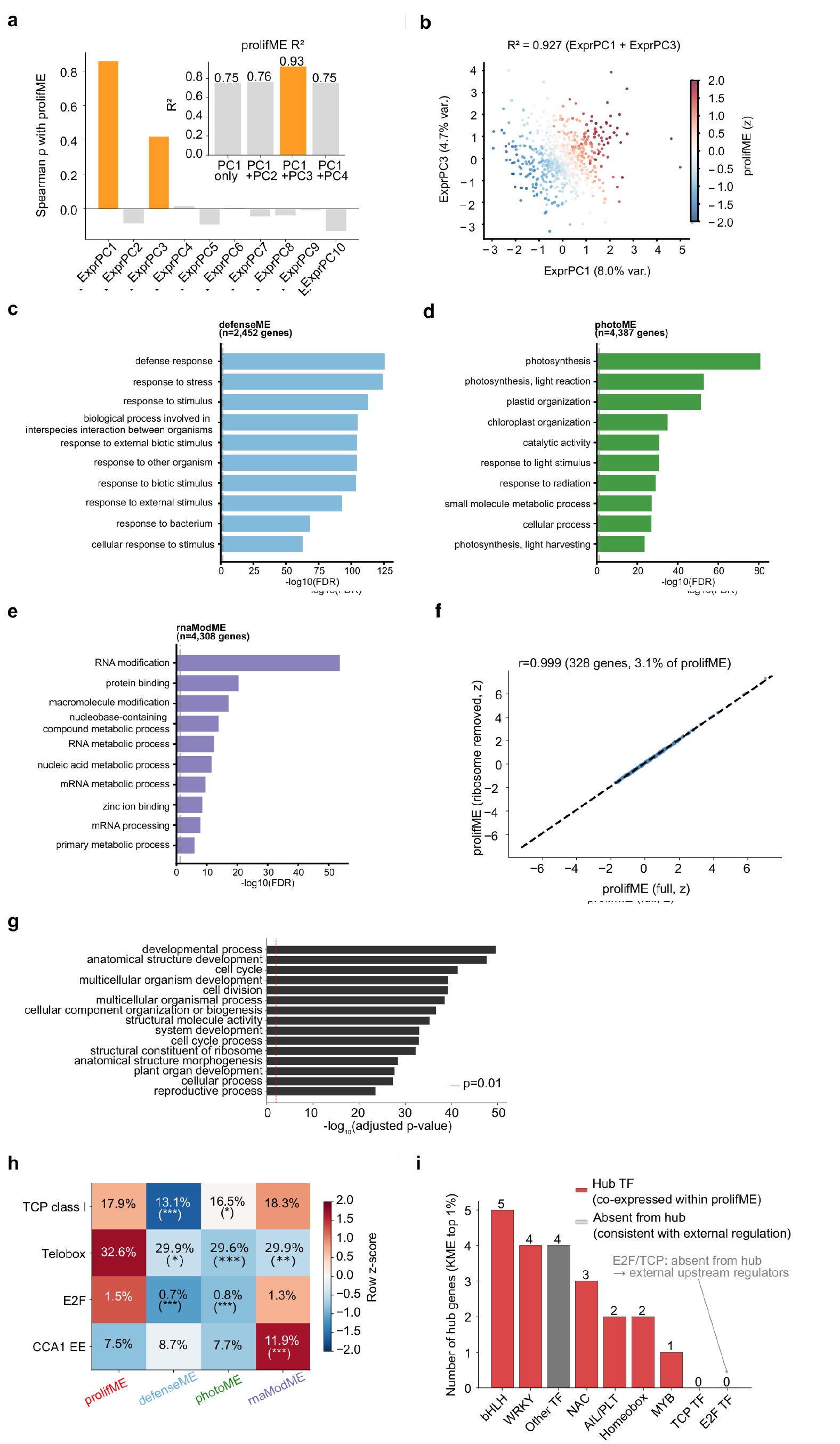
Identity and robustness of the four co-expression modules. (a) The prolifME eigengene against ExprPC1 (ρ = 0.858). (b) Variance in the prolifME eigengene explained by ExprPC1 and ExprPC3 together (92.7%). (c–e) Functional identity of the three comparator modules: defenseME, photoME and rnaModME. (f) The prolifME eigengene before and after removal of the 328 ribosomal genes (r = 0.999), showing that ribosomal abundance alone does not define the module. (g) The full GO biological-process enrichment for prolifME, which retains broad developmental terms; among the more specific terms cell-cycle regulation and ribosome biogenesis are the most prominent, and stress and environmental-sensing terms are not enriched. (h) Promoter motif enrichment for the cell-cycle and ribosome gene sets of all four modules, showing that the contrast in Fig. 2c is not specific to prolifME. (i) Top hub genes of prolifME (KME in the top 1%, 107 genes); neither the E2F nor the TCP class I factor families appear among them, which places both upstream of the program rather than at the center of its network.

**Fig. S4.**
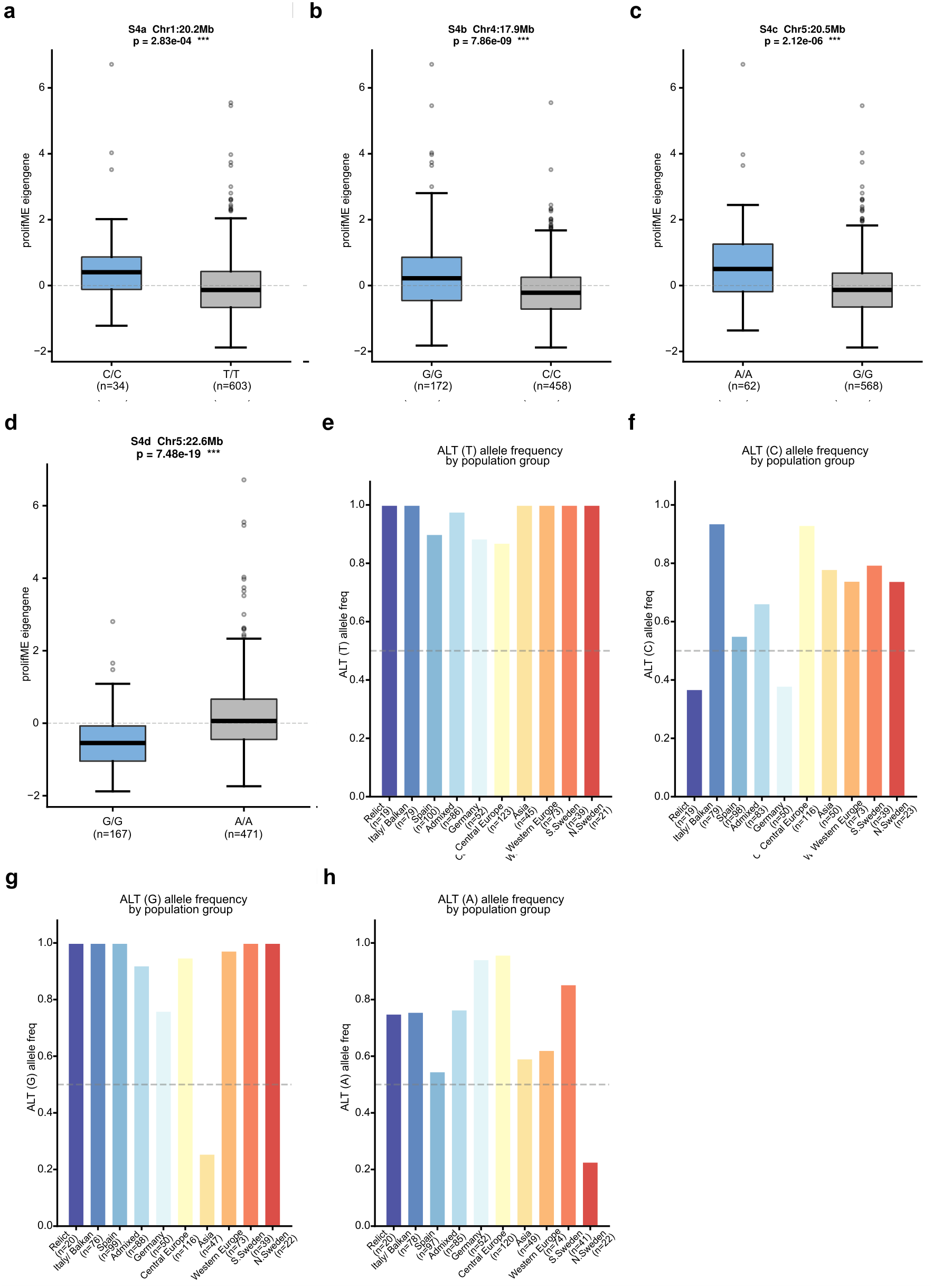
Genotype effects and allele frequencies at the four associated loci. (a–d) The prolifME eigengene by genotype at Chr1:20.2 Mb, Chr4:17.9 Mb, Chr5:20.5 Mb and Chr5:22.6 Mb, with the two-sided test P for each locus. Because Arabidopsis thaliana accessions are predominantly homozygous, only homozygous genotype classes are shown. (e–h) ALT allele frequency at the same four loci across the 1001 Genomes admixture groups, ordered by mean latitude and reported separately for southern and northern Sweden. At Chr5:22.6 Mb the allele associated with higher prolifME reaches about 0.75 in the relict and Spanish groups and is least frequent, about 0.22, in northern Sweden.

**Fig. S5.**
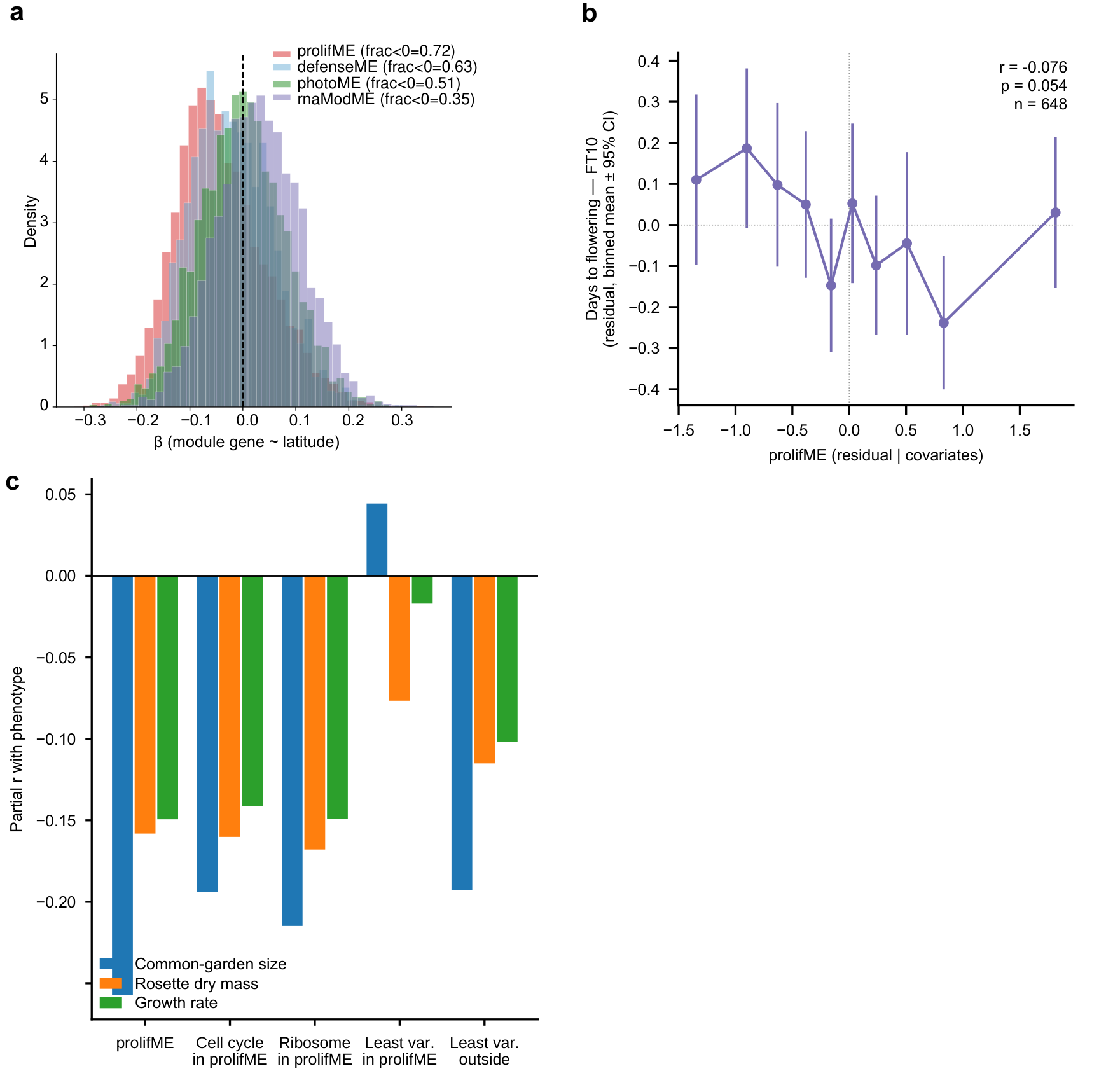
Directional coherence, flowering time and benchmarking against matched control gene sets. (a) Fraction of genes whose expression decreases with latitude, for each of the four modules: 72.0% for prolifME, 62.7% for defenseME, 50.7% for photoME and 35.4% for rnaModME. Significance is assessed against matched gene sets rather than a binomial null (Methods S1). (b) The prolifME eigengene against flowering time at 10 °C (r = −0.076, P = 0.054, n = 648), which is in the expected direction but not significant; the association strengthens to r = −0.131 (P = 8.3 × 10⁻⁴) when the developmental-stage covariate is added (Table S6). Correlations are partial on genomicPC1–10, latitude, longitude and ClockPC1. (c) Partial correlation with each of the three growth-strategy phenotypes — common-garden size, rosette dry mass and growth rate — for prolifME and for four matched control sets: cell-cycle and ribosomal subsets drawn from inside the module, and the least variable genes inside and outside it. The cell-cycle and ribosomal subsets track the module (r = −0.14 to −0.21) whereas genes drawn from inside it with low variance do not (r = +0.04 to −0.08). A size-matched random module is not an informative null at this module size, because random sets reach their asymptotic correlation by about 1,000 genes; significance was therefore assessed by permuting the phenotype labels with the gene set held fixed, and prolifME exceeded that null for the three traits shown here (P = 0.023–0.039). The fourth reported trait, δ¹³C, is not among the composition controls.

**Fig. S6.**
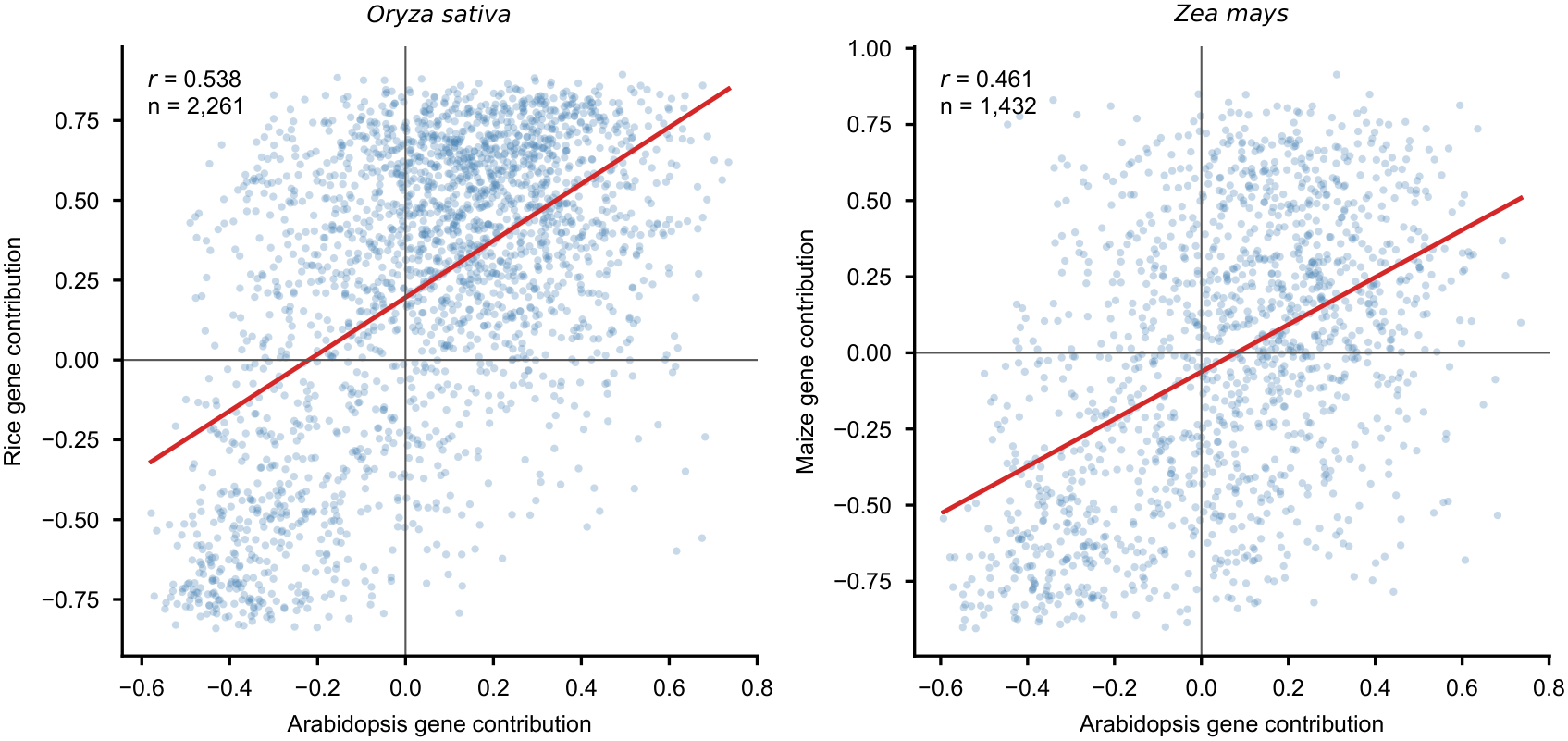
Cross-species alignment restricted to strict one-to-one orthologs. Arabidopsis gene contributions against contributions estimated independently in each species, for orthogroups with exactly one member per species: r = 0.538 in rice (n = 2,261) and r = 0.461 in maize (n = 1,432). Both exceed the whole-orthogroup values in Fig. 4a (0.446 and 0.379), indicating that ortholog ambiguity weakens rather than creates the alignment.

**Fig. S7.**
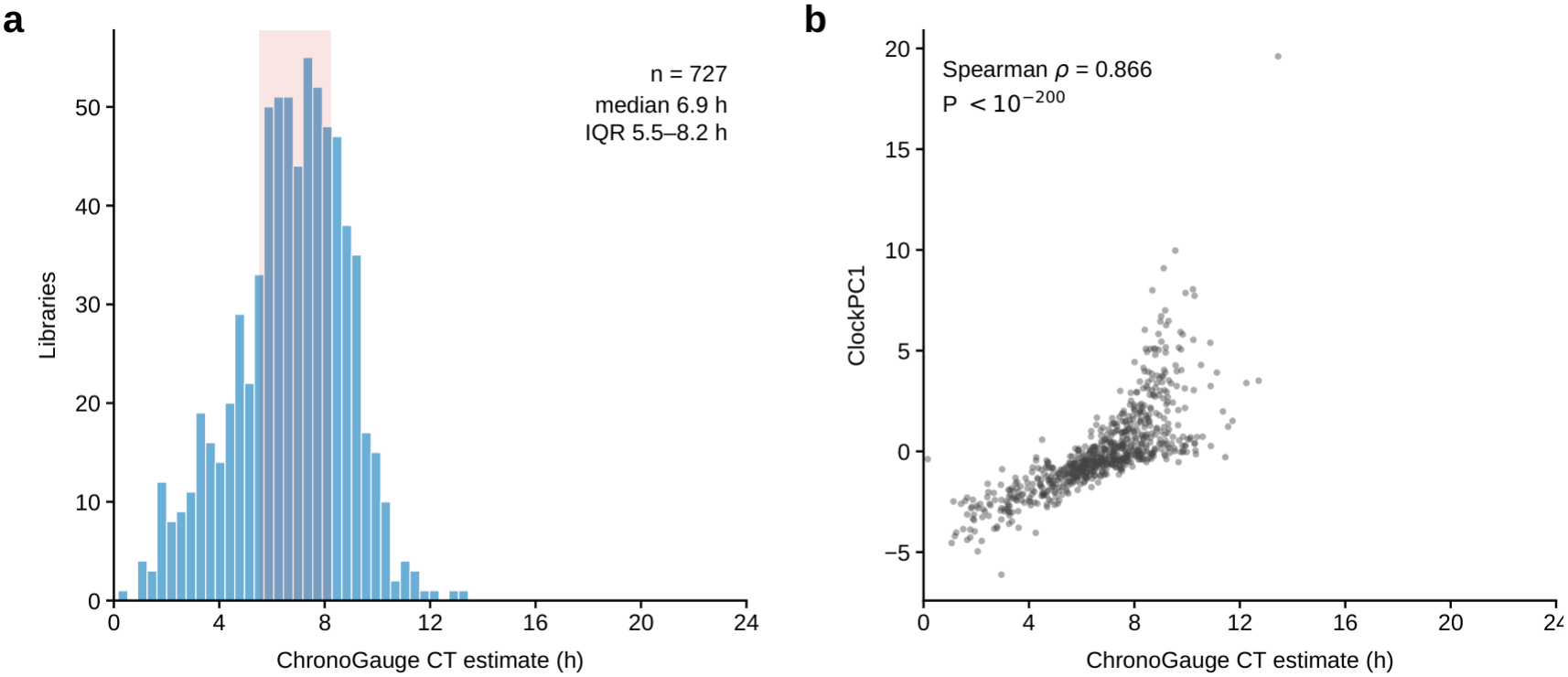
An independent circadian-time predictor relates ClockPC1 to predicted circadian state. ChronoGauge (Reynolds et al. 2025) was applied to all 727 libraries of the expression matrix using 84 of its 100 released sub-predictors; the remaining 16 require two genes that are absent from this matrix. Transcripts per million were reconstructed from the upper-quartile-normalized counts, which is exact because that normalization applies one scale factor per library and transcripts per million are a within-library composition. (a) Distribution of the ensemble circadian-time estimate across libraries; the shaded band marks the interquartile range. (b) The same estimate against ClockPC1 computed on these libraries by the procedure used throughout (Spearman ρ = 0.866, P < 1 × 10⁻²⁰⁰, n = 727). On the test set released with ChronoGauge this reimplementation reached a mean absolute error of 44 min (n = 58).

Table S1 Candidate genes at associated loci and their module membership scores.

Table S2 Variance explained by lead SNPs.

Table S3 Variance partitioning of ExprPC1 under four allocations.

Table S4 Per-axis variance partitioning for ExprPC1–10.

Table S5 Within-sample and cross-validated variance explained by selected SNPs.

Table S6 Phenotype associations with the developmental-stage covariate added.

Table S7 Four-block variance partitioning with the alternative circadian axis.

Table S8 Leave-one-group-out variance partitioning by admixture group.

Table S9 Sensitivity of module membership and eigengene to the clustering cut height.

Table S10 Cross-species alignment restricted to each module’s own orthogroups.

Table S11 SNP heritability of module eigengenes and expression axes.

Table S12 Transcriptome-wide variance partitioning by redundancy analysis.

Table S13 Screen of 461 AraPheno phenotypes with n ≥ 30, drawn from 535 phenotype files, with Benjamini–Hochberg correction.

