## supplementary tables for "Ancestry predicts a heritable proliferation state in *Arabidopsis*"

**Table S1** Candidate genes at associated loci and their module membership scores. KME is the Pearson correlation between each gene's expression and the prolifME eigengene across 664 accessions; genes with KME in the top 1% of the module (KME ≥ 0.683) are marked as hub genes.

| Locus | Gene | Symbol | Module | KME | Hub (top 1%) |
|---|---|---|---|---|---|
| Chr1:20.2 Mb | AT1G55580 | GRAS TF | prolifME | 0.667 | no |
| Chr1:20.2 Mb | AT1G54160 | NF-YA5 | prolifME | -0.1 | no |
| Chr4:17.9 Mb | AT4G37740 | GRF2 | prolifME | 0.448 | no |
| Chr4:17.9 Mb | AT4G38210 | EXP A20 | prolifME | 0.598 | no |
| Chr5:20.5 Mb | AT5G50820 | NAC97 | prolifME | 0.512 | no |
| Chr5:20.5 Mb | AT5G48560 | bHLH | prolifME | 0.561 | no |
| Chr5:22.6 Mb | AT5G56940 | RPS16 | rnaModME | 0.207 | no |
| Chr5:22.6 Mb | AT5G58290 | RPT3 | prolifME | 0.573 | no |

**Table S2** Variance explained by lead SNPs. $\Delta R^2$ is the gain in $R^2$ for each locus over genomicPC1–10 fitted alone, so the four values are marginal and are not additive; the four loci fitted jointly, with correction for selection, are in Table S5. Variant identifiers are absent from the genotype file and the loci are named by position.

| Lead SNP | Counted allele | n accessions | $\Delta R^2$ over genomicPC1–10 (%) | Model $R^2$ (%) | P | Genotype source |
|---|---|---|---|---|---|---|
| Chr1:20.2 Mb | C | 637 | 4.35 | 19.4 | 1.02e-08 | ld1_snp.raw |
| Chr4:17.9 Mb | G | 630 | 5.03 | 19.86 | 8.66e-10 | ld4_snp.raw |
| Chr5:20.5 Mb | A | 630 | 4.01 | 18.66 | 5.07e-08 | ld5a_snp.raw |
| Chr5:22.6 Mb | A | 638 | 7.25 | 22.2 | 8.21e-14 | ld5b_snp.raw |

**Table S3** Variance partitioning of ExprPC1 under four allocations.

| Predictor allocation | Genomic structure (%) | ClockPC1 (%) | Climate and soil (%) | Shared (%) | Residual (%) | Genomic / environmental |
|---|---|---|---|---|---|---|
| As described (coordinates environmental) | 4.98 | 19.78 | 2.63 | 7.42 | 65.19 | 1.9 |
| Coordinates with the genetic block | 8.04 | 19.53 | 2.25 | 5.01 | 65.18 | 3.57 |
| Coordinates excluded | 7.47 | 20.19 | 1.87 | 4.52 | 65.95 | 3.99 |
| Genomic components only (ClockPC1 and coordinates excluded) | 8.76 | | 1.41 | | | 6.21 |
| Two-block (genomic structure and ClockPC1 combined) | 26.05 | — | 2.63 | 6.13 | 65.19 | — |

**Table S4** Per-axis variance partitioning for ExprPC1–10. The structural block is genomicPC1–10, latitude, longitude and ClockPC1; the environmental block is the 19 bioclimatic and eight soil variables.

| Expression PC | PC variance (% of transcriptome) | Unique structural (% of PC) | Unique environmental (% of PC) | Shared (% of PC) | Residual (% of PC) | Unique structural (% of transcriptome) | Unique environmental (% of transcriptome) |
|---|---|---|---|---|---|---|---|
| ExprPC1 | 8.011 | 29.58 | 2.25 | 2.99 | 65.18 | 2.37 | 0.18 |
| ExprPC2 | 5.44 | 25.1 | 1.94 | 0.96 | 72 | 1.365 | 0.105 |
| ExprPC3 | 4.682 | 25.19 | 2.04 | 0.66 | 72.12 | 1.179 | 0.095 |
| ExprPC4 | 4.268 | 15.7 | 1.07 | 13.51 | 69.72 | 0.67 | 0.046 |
| ExprPC5 | 3.264 | 12.62 | 1.27 | 2.74 | 83.37 | 0.412 | 0.042 |
| ExprPC6 | 3.122 | 4.45 | 2.88 | 1 | 91.67 | 0.139 | 0.09 |
| ExprPC7 | 2.611 | 5.27 | 1.96 | 7.47 | 85.3 | 0.138 | 0.051 |
| ExprPC8 | 2.148 | 8.03 | 2.59 | 6.82 | 82.56 | 0.173 | 0.056 |
| ExprPC9 | 1.986 | 7.38 | 2.95 | 2.4 | 87.27 | 0.147 | 0.059 |
| ExprPC10 | 1.648 | 10.32 | 2.31 | 1.37 | 86 | 0.17 | 0.038 |

**Table S5** Within-sample and cross-validated variance explained by selected SNPs.

| Estimate | Value (% of eigengene variance) |
|---|---|
| Within-sample adjusted $R^2$ (as reported) | 15.45 |
| Permutation null — $R^2$ from selection alone | 11.49 |
| Observed minus null | 3.96 |
| Cross-validated (SNP set fixed) | 13.57 |
| Cross-validated, selection repeated per fold | 2.68 |

**Table S6** Phenotype associations with the developmental-stage covariate added.

| Section | Row | Env. unique (%) | genomicPC (%) | ClockPC1 (%) | StagePC1 (%) | Shared (%) | Unexplained (%) | n | r | P | r (declared) | P (declared) | r (+StagePC1) | P (+StagePC1) | Attenuation | Verdict |
|---|---|---|---|---|---|---|---|---|---|---|---|---|---|---|---|---|
| Four-block partition | ExprPC1 | 3.13 | 5.96 | 19.06 | 6.47 | 6.65 | 58.73 | 665 | | | | | | | | |
| Four-block partition | prolifME | 3.85 | 5.18 | 4.37 | 1.89 | 9.73 | 74.98 | 664 | | | | | | | | |
| StagePC1 external | ClockPC1 | | | | | | | 665 | 0.006 | 0.869 | | | | | | |
| StagePC1 external | ExprPC1 | | | | | | | 665 | -0.214 | 2.43e-08 | | | | | | |
| StagePC1 external | ExprPC3 | | | | | | | 665 | 0.026 | 0.509 | | | | | | |
| StagePC1 external | prolifME | | | | | | | 664 | -0.131 | 0.000721 | | | | | | |
| StagePC1 external | lat_x | | | | | | | 664 | 0.085 | 0.0286 | | | | | | |
| StagePC1 external | FT10 | | | | | | | 650 | -0.378 | 1.89e-23 | | | | | | |
| Trait association | SizeLocSweden | | | | | | | 80 | | | -0.2576 | 0.021 | -0.3163 | 0.0043 | -0.2275 | Survives |
| Trait association | rosetteDM | | | | | | | 253 | | | -0.1584 | 0.0117 | -0.1635 | 0.0092 | -0.0327 | Survives |
| Trait association | GrowthRate | | | | | | | 253 | | | -0.1496 | 0.0173 | -0.1563 | 0.0128 | -0.0452 | Survives |
| Trait association | delta13C | | | | | | | 149 | | | -0.1987 | 0.0151 | -0.1881 | 0.0216 | 0.0532 | Survives |
| Trait association | FT10 | | | | | | | 648 | | | -0.0759 | 0.0535 | -0.1309 | 0.0008 | -0.7255 | Arises only with the covariate |

**Table S7** Four-block variance partitioning with the alternative circadian axis.

| Response | Base model | ΔR² genomic | p (permutation) | ΔR² climate and soil | Base model R² | ΔR² added by CircPC1 over ClockPC1 | Ordering holds |
|---|---|---|---|---|---|---|---|
| ExprPC1 | ClockPC1 only | 0.1165 | 0.005 | 0.0592 | 0.265 | 0.2823 | yes |
| ExprPC1 | CircPC1 only | 0.057 | 0.005 | 0.0253 | 0.5398 | 0.2823 | yes |
| ExprPC1 | Both | 0.0537 | 0.005 | 0.0228 | 0.5472 | 0.2823 | yes |
| ExprPC1 | Sensitivity: coordinates in environmental block | 0.0537 | 0.005 | 0.0465 | 0.5472 | 0.2823 | yes |
| ExprPC1 | Two-block allocation D, as defined in the text | 0.0876 | | 0.0141 | 0.051 | 0.2823 | yes |
| ExprPC1 | Mutually conditioned on circadian and stage axes | 0.0389 | | 0.008 | 0.5472 | 0.2823 | yes |
| prolifME | ClockPC1 only | 0.1374 | 0.005 | 0.0556 | 0.0744 | 0.0702 | yes |
| prolifME | CircPC1 only | 0.1129 | 0.005 | 0.0439 | 0.143 | 0.0702 | yes |
| prolifME | Both | 0.1117 | 0.005 | 0.0431 | 0.1445 | 0.0702 | yes |
| prolifME | Sensitivity: coordinates in environmental block | 0.1117 | 0.005 | 0.1076 | 0.1445 | 0.0702 | yes |
| prolifME | Two-block allocation D, as defined in the text | 0.1042 | | 0.0127 | 0.0556 | 0.0702 | yes |
| prolifME | Mutually conditioned on circadian and stage axes | 0.0799 | | 0.0112 | 0.1445 | 0.0702 | yes |
| ExprPC1 | Four-block ancestry unique (ClockPC1) | 0.0596 | | | | | |
| ExprPC1 | Four-block ancestry unique (CircPC1 substituted) | 0.0239 | | | | | |
| prolifME | Four-block ancestry unique (ClockPC1) | 0.0518 | | | | | |
| prolifME | Four-block ancestry unique (CircPC1 substituted) | 0.0374 | | | | | |

**Table S8** Leave-one-group-out variance partitioning by admixture group.

| Response | All accessions (%) | Relict accessions removed (%) | Leave-one-group-out min (%) | Leave-one-group-out median (%) | Leave-one-group-out max (%) |
|---|---|---|---|---|---|
| ExprPC1 | 8.76 | 9.55 | 6.81 | 9.23 | 9.9 |
| prolifME | 10.42 | 10.73 | 7.21 | 10.82 | 11.56 |

**Table S9** Sensitivity of module membership and eigengene to the clustering cut height.

| Cut height | Modules | Unassigned genes | Proliferative module size | Jaccard vs published module | \|r\| with published eigengene |
|---|---|---|---|---|---|
| 0.93 | 38 | 2614 | 5468 | 0.5093 | 0.9568 |
| 0.95 | 36 | 1890 | 5620 | 0.5235 | 0.9572 |
| 0.96 | 34 | 1272 | 5730 | 0.5337 | 0.9571 |
| 0.97 | 29 | 838 | 5763 | 0.5368 | 0.9571 |
| 0.98 | 23 | 335 | 10736 | 1 | 1 |
| 0.99 | 11 | 26 | 15466 | 0.6942 | 0.9801 |

**Table S10** Cross-species alignment restricted to each module's own orthogroups.

| Comparison | Species | Module | r | n orthogroups |
|---|---|---|---|---|
| All shared orthogroups | rice | photoME | -0.471 | 10182 |
| All shared orthogroups | rice | prolifME | 0.446 | 10182 |
| All shared orthogroups | rice | rnaModME | 0.163 | 10182 |
| All shared orthogroups | rice | defenseME | -0.085 | 10182 |
| All shared orthogroups | maize | prolifME | 0.379 | 10903 |
| All shared orthogroups | maize | photoME | -0.306 | 10903 |
| All shared orthogroups | maize | defenseME | -0.112 | 10903 |
| All shared orthogroups | maize | rnaModME | 0.092 | 10903 |
| Module's own orthogroups | rice | photoME | -0.576 | 2141 |
| Module's own orthogroups | rice | prolifME | 0.046 | 4624 |
| Module's own orthogroups | rice | rnaModME | -0.019 | 2208 |
| Module's own orthogroups | rice | defenseME | -0.179 | 776 |
| Module's own orthogroups | maize | prolifME | 0.107 | 4955 |
| Module's own orthogroups | maize | photoME | -0.43 | 2272 |
| Module's own orthogroups | maize | defenseME | -0.185 | 814 |
| Module's own orthogroups | maize | rnaModME | -0.057 | 2329 |
| Paired bootstrap (prolifME vs best other) | rice | top in 0.000 of resamples | -0.025 | 10182 |
| Paired bootstrap (prolifME vs best other) | maize | top in 1.000 of resamples | 0.074 | 10903 |

**Table S11** SNP heritability of module eigengenes and expression axes.

| Trait | n | SNP h2 | 95% profile interval | p (LRT) | h2 \| genomicPC1-10 fixed | Adjusted R2 on genomicPC1-10 |
|---|---|---|---|---|---|---|
| ClockPC1 | 665 | 0.0232 | 0.00–0.13 | 0.1489 | 0.0312 | 0.0037 |
| prolifME | 664 | 0.1925 | 0.12–0.29 | 0 | 0.1684 | 0.1341 |
| defenseME | 664 | 0.2616 | 0.15–0.40 | 0 | 0.2508 | 0.0718 |
| photoME | 664 | 0.2147 | 0.09–0.37 | 0 | 0.2694 | 0.0597 |
| rnaModME | 664 | 0.3026 | 0.12–0.52 | 0 | 0.302 | 0.0523 |
| ExprPC1 | 665 | 0.2084 | 0.12–0.32 | 0 | 0.2039 | 0.1099 |
| ExprPC2 | 665 | 0.2546 | 0.10–0.44 | 0 | 0.3065 | 0.0495 |
| ExprPC3 | 665 | 0.4253 | 0.25–0.62 | 0 | 0.4296 | 0.0942 |
| ExprPC4 | 665 | 0.4188 | 0.27–0.58 | 0 | 0.3465 | 0.191 |
| ExprPC5 | 665 | 0.2521 | 0.12–0.42 | 0 | 0.2732 | 0.0826 |

**Table S12** Transcriptome-wide variance partitioning by redundancy analysis.

| Allocation | Structural unique (%) | Environmental unique (%) | Shared (%) | Both blocks (%) | Structural alone (%) | Environmental alone (%) | Unexplained (%) |
|---|---|---|---|---|---|---|---|
| A as-declared (coords→env) | 6.1 | 2.19 | 3.58 | 11.88 | 9.69 | 5.78 | 88.12 |
| B coords→structural | 6.68 | 1.89 | 3.31 | 11.88 | 9.99 | 5.2 | 88.12 |
| C coords excluded | 6.5 | 2.02 | 3.19 | 11.7 | 9.69 | 5.2 | 88.3 |
| D genomic only (no clock) | 2.55 | 2 | 3.2 | 7.75 | 5.75 | 5.2 | 92.25 |
| Three-block: genomicPC1-10 unique | 2.46 | | | | | | |
| Three-block: Environmental unique | 2.02 | | | | | | |
| Three-block: ClockPC1 unique | 3.95 | | | | | | |
| Three-block: Shared | 3.27 | | | | | | |
| Three-block: All three blocks | 11.71 | | | | | | |
| Three-block: Unexplained | 88.3 | | | | | | |
| Per axis (STRUCT vs ENV): ExprPC1 | 21.12 | 5.7 | | | | | |
| Per axis (STRUCT vs ENV): ExprPC2 | 28.54 | 2.66 | | | | | |
| Per axis (STRUCT vs ENV): ExprPC3 | 12.31 | 2.96 | | | | | |
| Per axis (STRUCT vs ENV): ExprPC4 | 24.01 | 0.08 | | | | | |
| Per axis (STRUCT vs ENV): ExprPC5 | 7.47 | 3.47 | | | | | |

**Table S13** Screen of 461 AraPheno phenotypes with n ≥ 30, drawn from 535 phenotype files, with Benjamini–Hochberg correction.

| **Table S13 Screen of the proliferation module eigengene against the AraPheno phenotype collection.** | | | | | | | |
|---|---|---|---|---|---|---|---|
| Each phenotype was correlated with the prolifME eigengene by simple Pearson correlation with no covariates. Of 535 phenotype files, 461 shared at least 30 accessions with the expression data and entered the screen; Benjamini-Hochberg correction was applied across those 461 tests. Because no covariates are included, the coefficients here are systematically larger than the partial correlations reported in the main text (rosette dry mass -0.244 here against -0.151 under the covariate model). This table documents how the reported traits were selected and is not the source of their effect sizes. Traits reported in the main text are flagged in the final column. | | | | | | | |
| Rank | Phenotype | AraPheno file | n accessions | Pearson r | P | q (BH) | Reported in main text |
| 1 | rosetteDM | 390.csv | 253 | -0.2439 | 8.86831e-05 | 0.0363 | Fig. 3d (rosette dry mass) |
| 2 | FT10 | 261.csv | 648 | -0.1479 | 0.000157508 | 0.0363 | Fig. S5b (flowering time, 10 °C) |
| 3 | Leaf epidermis autofluorescence | 1900.csv | 120 | 0.3124 | 0.000512197 | 0.0687 | |
| 4 | GrowthRate | 391.csv | 253 | -0.2144 | 0.000596421 | 0.0687 | Fig. 3e (growth rate) |
| 5 | As2 CFU2 | 21.csv | 46 | 0.4351 | 0.0025143 | 0.2021 | |
| 6 | SizeLocSweden2009 | 115.csv | 80 | -0.3285 | 0.00293144 | 0.2021 | Fig. 3c (common-garden size) |
| 7 | ScalingExponent | 392.csv | 253 | 0.1851 | 0.00311688 | 0.2021 | |
| 8 | delta 13C 261 | 753.csv | 149 | -0.232 | 0.00441367 | 0.2021 | Fig. 3f (δ13C) |
| 9 | RL | 704.csv | 573 | -0.1188 | 0.0044174 | 0.2021 | |
| 10 | FT16 | 262.csv | 630 | -0.112 | 0.00487885 | 0.2021 | |
| 11 | As2 | 103.csv | 46 | 0.4074 | 0.00494785 | 0.2021 | |
| 12 | delta 13C | 752.csv | 179 | -0.2062 | 0.00561754 | 0.2021 | (same trait name, not the reported file: Fig. 3f (δ13C)) |
| 13 | Size sweden 2009 (1st experiment) | 123.csv | 80 | -0.3064 | 0.00569948 | 0.2021 | |
| 14 | CDT (Controlled seed Deterioration Treatment) | 1309.csv | 40 | 0.4208 | 0.00685231 | 0.207 | |
| 15 | CL | 705.csv | 603 | -0.1095 | 0.00711826 | 0.207 | |
| 16 | DTF1 | 703.csv | 619 | -0.108 | 0.00718293 | 0.207 | |
| 17 | Trichome stem length | 1898.csv | 118 | -0.242 | 0.00829272 | 0.2222 | |
| 18 | DTFplantingSummer2009 | 129.csv | 83 | -0.2847 | 0.00909534 | 0.2222 | |
| 19 | DTF2 | 701.csv | 617 | -0.1048 | 0.00915982 | 0.2222 | |
| 20 | DTFlocSweden2009 | 118.csv | 83 | -0.2822 | 0.00974384 | 0.2246 | |
| 21 | At2 CFU2 | 2.csv | 46 | 0.3708 | 0.0111903 | 0.2457 | |
| 22 | DTF3 | 702.csv | 612 | -0.1007 | 0.0127229 | 0.2666 | |
| 23 | DTF sweden 2009 (2nd experiment) | 140.csv | 83 | -0.2648 | 0.0155632 | 0.3119 | |
| 24 | Size sweden 2009 (2nd experiment) | 110.csv | 75 | -0.2739 | 0.0174158 | 0.3345 | |
| 25 | M172T666 | 146.csv | 107 | 0.2212 | 0.0220542 | 0.4067 | |
| 26 | DTF spain 2009 (2nd experiment) | 121.csv | 83 | -0.2463 | 0.0248023 | 0.4357 | |
| 27 | DTF sweden 2009 (1st experiment) | 134.csv | 83 | -0.2451 | 0.0255163 | 0.4357 | |
| 28 | M130T666 | 145.csv | 107 | 0.21 | 0.0299626 | 0.4933 | |

| **Table S13 Screen of the proliferation module eigengene against the AraPheno phenotype collection.** | | | | | | | |
|---|---|---|---|---|---|---|---|
| 29 | FT10 | 31.csv | 53 | -0.2737 | 0.0473917 | 0.6936 | (same trait name, not the reported file: Fig. S5b (flowering time, 10 °C)) |
| 30 | rSeeds tlp | 541.csv | 274 | 0.1195 | 0.048053 | 0.6936 | |
| 31 | SizePlantingSummerLocSweden2009 | 111.csv | 75 | 0.2289 | 0.0482152 | 0.6936 | |
| 32 | Trichome circularity | 1893.csv | 116 | 0.183 | 0.0492115 | 0.6936 | |
| 33 | LD | 89.csv | 46 | -0.2896 | 0.0509191 | 0.6936 | |
| 34 | Raffinose 32C | 1910.csv | 131 | -0.1708 | 0.0511548 | 0.6936 | |
| 35 | rFitness mhi | 518.csv | 337 | 0.1024 | 0.0604561 | 0.7515 | |
| 36 | LN16 | 87.csv | 45 | -0.2807 | 0.0617521 | 0.7515 | |
| 37 | rSeeds thp | 539.csv | 337 | 0.1015 | 0.0626487 | 0.7515 | |
| 38 | FT16 | 30.csv | 52 | -0.2592 | 0.0634926 | 0.7515 | |
| 39 | rFitness thp | 523.csv | 337 | 0.1 | 0.0668153 | 0.7515 | |
| 40 | root length day001 minusZn | 761.csv | 40 | 0.2927 | 0.0668327 | 0.7515 | |
| 41 | Z-mean Total length day003 | 745.csv | 40 | 0.2927 | 0.0668327 | 0.7515 | |
| 42 | rSurvival fruit tli | 532.csv | 337 | 0.0965 | 0.076966 | 0.8448 | |
| 43 | M216T665 | 144.csv | 107 | 0.1703 | 0.079399 | 0.8512 | |
| 44 | FT22 | 102.csv | 53 | -0.2397 | 0.0838813 | 0.8624 | |
| 45 | Trichome perimeter | 1896.csv | 116 | -0.161 | 0.084186 | 0.8624 | |
| 46 | Dolichole | 1312.csv | 34 | -0.2967 | 0.0883839 | 0.8777 | |
| 47 | Trichome solidity | 1894.csv | 116 | 0.1578 | 0.0906409 | 0.8777 | |
| 48 | Trichome length | 1895.csv | 116 | -0.1555 | 0.09548 | 0.8777 | |
| 49 | rSeeds thi | 538.csv | 337 | 0.0906 | 0.0969181 | 0.8777 | |
| 50 | root length day003 minusZn | 763.csv | 47 | 0.2434 | 0.0991844 | 0.8777 | |
| 51 | Z-mean Total length day005 | 747.csv | 47 | 0.2434 | 0.0991844 | 0.8777 | |
| 52 | Se82 | 284.csv | 58 | -0.2183 | 0.0996714 | 0.8777 | |
| 53 | clim-bio8 | 571.csv | 327 | -0.0908 | 0.101289 | 0.8777 | |
| 54 | Carotenoid | 1313.csv | 35 | 0.2804 | 0.102806 | 0.8777 | |
| 55 | DTFlocSweden2008 | 117.csv | 59 | -0.2124 | 0.106247 | 0.8842 | |
| 56 | FT GH | 49.csv | 46 | -0.2382 | 0.110935 | 0.8842 | |
| 57 | Germ 22 | 45.csv | 46 | -0.2372 | 0.112387 | 0.8842 | |
| 58 | LRDpBZ75 | 667.csv | 51 | -0.2237 | 0.114608 | 0.8842 | |
| 59 | Germ 16 | 60.csv | 45 | -0.2384 | 0.114845 | 0.8842 | |
| 60 | Raffinose 22C | 1911.csv | 131 | -0.1379 | 0.116273 | 0.8842 | |
| 61 | stomata density | 748.csv | 190 | -0.1141 | 0.116998 | 0.8842 | |
| 62 | Width 22 | 26.csv | 46 | 0.2325 | 0.119899 | 0.8915 | |
| 63 | Metal gradient in trichomes | 1905.csv | 119 | -0.1418 | 0.124081 | 0.908 | |
| 64 | Chlorophyll | 1311.csv | 35 | 0.2632 | 0.126645 | 0.9122 | |
| 65 | At1 | 84.csv | 46 | 0.2266 | 0.129965 | 0.9161 | |
| 66 | Bs | 10.csv | 46 | 0.2253 | 0.13229 | 0.9161 | |
| 67 | Basal125 | 691.csv | 51 | -0.2132 | 0.133149 | 0.9161 | |
| 68 | rFitness thi | 522.csv | 337 | 0.0814 | 0.136042 | 0.9164 | |
| 69 | LN10 | 85.csv | 46 | -0.2217 | 0.138739 | 0.9164 | |
| 70 | Size spain 2009 (2nd experiment) | 127.csv | 76 | -0.1712 | 0.139152 | 0.9164 | |

| **Table S13 Screen of the proliferation module eigengene against the AraPheno phenotype collection.** | | | | | | |
|---|---|---|---|---|---|---|
| 71 | rSeeds mhi | 534.csv | 336 | 0.0794 | 0.146604 | 0.9256 |
| 72 | greening 15d protocol a | 1273.csv | 36 | -0.2468 | 0.146775 | 0.9256 |
| 73 | FruitNumber | 388.csv | 247 | 0.0925 | 0.14735 | 0.9256 |
| 74 | root length day005 minusZn | 765.csv | 46 | 0.2159 | 0.149645 | 0.9256 |
| 75 | Mean(LRD P) C | 356.csv | 30 | -0.269 | 0.150587 | 0.9256 |
| 76 | rSurvival fruit thp | 531.csv | 337 | 0.0769 | 0.158812 | 0.9608 |
| 77 | RGR | 389.csv | 249 | 0.089 | 0.161546 | 0.9608 |
| 78 | Z-mean Total length day004 | 746.csv | 48 | 0.2019 | 0.168671 | 0.9608 |
| 79 | root length day002 minusZn | 762.csv | 48 | 0.2019 | 0.168671 | 0.9608 |
| 80 | Gravitropic setpoint angle (GSA) | 768.csv | 52 | -0.1932 | 0.169948 | 0.9608 |
| 81 | clim-bio7 | 570.csv | 327 | 0.0759 | 0.171082 | 0.9608 |
| 82 | clim-tmin12 | 631.csv | 316 | -0.0768 | 0.172978 | 0.9608 |
| 83 | Callose content in trichomes | 1897.csv | 116 | 0.1274 | 0.172991 | 0.9608 |
| 84 | LN22 | 41.csv | 46 | -0.2032 | 0.175664 | 0.9641 |
| 85 | clim-bio4 | 567.csv | 327 | 0.0741 | 0.181621 | 0.9747 |
| 86 | DTFmainEffect2009 | 125.csv | 83 | -0.148 | 0.181832 | 0.9747 |
| 87 | DTF spain 2009 (1st experiment) | 119.csv | 83 | -0.1463 | 0.186773 | 0.9761 |
| 88 | root length day007 minusZn | 767.csv | 48 | 0.1924 | 0.190109 | 0.9761 |
| 89 | Length | 700.csv | 457 | 0.0613 | 0.190706 | 0.9761 |
| 90 | greening 15d protocol b | 1280.csv | 41 | -0.2074 | 0.193271 | 0.9761 |
| 91 | clim-tmax5 | 623.csv | 327 | 0.0719 | 0.194668 | 0.9761 |
| 92 | Mean(P2) ABA | 373.csv | 30 | 0.2412 | 0.199175 | 0.9761 |
| 93 | Number of trichome branches | 1903.csv | 119 | 0.1178 | 0.201915 | 0.9761 |
| 94 | FT Diameter Field | 1.csv | 48 | 0.1873 | 0.202384 | 0.9761 |
| 95 | PF-mean Total length day003 | 739.csv | 45 | 0.1936 | 0.20269 | 0.9761 |
| 96 | AAT (Accelerated seed Aging Treatment) | 1308.csv | 41 | 0.2028 | 0.203527 | 0.9761 |
| 97 | rSurvival fruit mhi | 526.csv | 337 | 0.0691 | 0.20558 | 0.9761 |
| 98 | DTF sweden 2008 (1st experiment) | 113.csv | 59 | -0.1655 | 0.210217 | 0.9761 |
| 99 | GR21 cold | 316.csv | 35 | -0.2145 | 0.215846 | 0.9761 |
| 100 | rFitness tli | 524.csv | 337 | 0.0672 | 0.21857 | 0.9761 |
| 101 | stomata density 261 | 749.csv | 149 | -0.1013 | 0.218861 | 0.9761 |
| 102 | Cu65 | 281.csv | 58 | 0.1638 | 0.219187 | 0.9761 |
| 103 | rSeeds tli | 540.csv | 329 | -0.0677 | 0.220823 | 0.9761 |
| 104 | 2W | 11.csv | 45 | -0.1845 | 0.225137 | 0.9761 |
| 105 | clim-aet10 | 543.csv | 327 | -0.0668 | 0.228671 | 0.9761 |
| 106 | Apical75 | 670.csv | 51 | -0.1711 | 0.230076 | 0.9761 |
| 107 | Anthocyanin 22 | 75.csv | 46 | -0.1797 | 0.231984 | 0.9761 |
| 108 | clim-cyclevar | 574.csv | 327 | 0.0661 | 0.233522 | 0.9761 |
| 109 | Area sweden 2009 (1st experiment) | 108.csv | 80 | -0.134 | 0.236079 | 0.9761 |
| 110 | Mean(P2) C | 355.csv | 30 | 0.2227 | 0.236942 | 0.9761 |
| 111 | Yield sweden 2009 (2nd experiment) | 114.csv | 55 | -0.1618 | 0.238063 | 0.9761 |
| 112 | 0W GH FT | 53.csv | 39 | -0.1923 | 0.240796 | 0.9761 |
| 113 | S34 | 274.csv | 58 | -0.1536 | 0.249648 | 0.9761 |
| 114 | SizePlantingSummer2009 | 109.csv | 76 | -0.1317 | 0.256791 | 0.9761 |
| 115 | LRL0 | 648.csv | 51 | 0.161 | 0.258929 | 0.9761 |

| **Table S13 Screen of the proliferation module eigengene against the AraPheno phenotype collection.** | | | | | | |
|---|---|---|---|---|---|---|
| 116 | root length day006 | 759.csv | 48 | 0.1652 | 0.261775 | 0.9761 |
| 117 | area 15d protocol b | 1279.csv | 41 | -0.1782 | 0.264931 | 0.9761 |
| 118 | DTFplantingSummer2008 | 130.csv | 59 | -0.1467 | 0.267395 | 0.9761 |
| 119 | MRVA0 | 643.csv | 51 | -0.1571 | 0.270848 | 0.9761 |
| 120 | Depth0 | 642.csv | 51 | 0.157 | 0.271102 | 0.9761 |
| 121 | SD | 47.csv | 40 | -0.1778 | 0.272301 | 0.9761 |
| 122 | RGR Zn | 721.csv | 48 | 0.1612 | 0.273695 | 0.9761 |
| 123 | Mean(LRD R) ABA | 375.csv | 30 | -0.2058 | 0.275205 | 0.9761 |
| 124 | clim-aet11 | 544.csv | 327 | -0.0604 | 0.276363 | 0.9761 |
| 125 | DTF sweden 2008 (2nd experiment) | 141.csv | 59 | -0.1429 | 0.280305 | 0.9761 |
| 126 | area 21d protocol b | 1293.csv | 41 | -0.1717 | 0.283202 | 0.9761 |
| 127 | clim-gs tmin | 591.csv | 327 | -0.0587 | 0.290178 | 0.9761 |
| 128 | clim-bio11 | 556.csv | 326 | -0.0584 | 0.293017 | 0.9761 |
| 129 | RGR Fe | 717.csv | 48 | 0.1544 | 0.294672 | 0.9761 |
| 130 | Leaf roll 10 | 9.csv | 46 | 0.1572 | 0.296866 | 0.9761 |
| 131 | 4W | 100.csv | 36 | -0.1779 | 0.299258 | 0.9761 |
| 132 | shoots 15d protocol a | 1267.csv | 36 | -0.1773 | 0.300817 | 0.9761 |
| 133 | F-mean Total length day005 | 735.csv | 48 | 0.1521 | 0.301975 | 0.9761 |
| 134 | FLC | 29.csv | 42 | -0.1625 | 0.30391 | 0.9761 |
| 135 | root length day004 | 757.csv | 48 | 0.1511 | 0.305192 | 0.9761 |
| 136 | clim-gs1 | 576.csv | 327 | -0.0568 | 0.305431 | 0.9761 |
| 137 | clim-tmax1 | 616.csv | 327 | -0.0568 | 0.305565 | 0.9761 |
| 138 | clim-tmax12 | 619.csv | 327 | -0.0568 | 0.305862 | 0.9761 |
| 139 | Trichome area | 1892.csv | 116 | -0.0954 | 0.30819 | 0.9761 |
| 140 | YieldPlantingSummer2009 | 133.csv | 67 | 0.1259 | 0.310115 | 0.9761 |
| 141 | clim-prec4 | 610.csv | 327 | -0.0562 | 0.311061 | 0.9761 |
| 142 | rSurvival fruit mli | 528.csv | 337 | 0.055 | 0.314092 | 0.9761 |
| 143 | Mean(LRR) C | 360.csv | 30 | -0.1899 | 0.314926 | 0.9761 |
| 144 | clim-tmin1 | 628.csv | 325 | -0.0557 | 0.316884 | 0.9761 |
| 145 | EPPO (Elevated Partial Pressure of Oxygen seed-aging treatment) | 1310.csv | 41 | 0.1596 | 0.318856 | 0.9761 |
| 146 | NAT (Natural seed Aging Treatment) | 1307.csv | 41 | 0.1565 | 0.328567 | 0.9761 |
| 147 | RGR MS | 718.csv | 48 | 0.1437 | 0.329923 | 0.9761 |
| 148 | Mean(TLRL) ABA | 376.csv | 30 | -0.183 | 0.33309 | 0.9761 |
| 149 | DTF spain 2008 (2nd experiment) | 136.csv | 59 | -0.1281 | 0.333472 | 0.9761 |
| 150 | F-mean Total length day004 | 734.csv | 48 | 0.1422 | 0.334869 | 0.9761 |
| 151 | clim-tmin10 | 629.csv | 325 | -0.0535 | 0.336601 | 0.9761 |
| 152 | YieldLocSweden2009 | 135.csv | 73 | -0.1136 | 0.338481 | 0.9761 |
| 153 | rSurvival fruit mlp | 529.csv | 337 | 0.0523 | 0.338664 | 0.9761 |
| 154 | DTFplantingSummerLocSweden2009 | 116.csv | 83 | 0.1062 | 0.339253 | 0.9761 |
| 155 | deltaRGR P Zn norm on Zn norm | 729.csv | 48 | -0.1408 | 0.339688 | 0.9761 |
| 156 | clim-bio6 | 569.csv | 324 | -0.053 | 0.341886 | 0.9761 |
| 157 | clim-tmax11 | 618.csv | 327 | -0.0527 | 0.341948 | 0.9761 |
| 158 | Trichome density | 1902.csv | 120 | -0.0871 | 0.34419 | 0.9761 |
| 159 | Mean(LR.no) C | 354.csv | 30 | -0.1788 | 0.344557 | 0.9761 |
| 160 | clim-bio2 | 565.csv | 327 | 0.0524 | 0.345105 | 0.9761 |

| **Table S13 Screen of the proliferation module eigengene against the AraPheno phenotype collection.** | | | | | | |
|---|---|---|---|---|---|---|
| 161 | SDV | 104.csv | 40 | -0.1515 | 0.350569 | 0.9761 |
| 162 | Mean(LRR) ABA | 378.csv | 30 | -0.1759 | 0.352474 | 0.9761 |
| 163 | As | 107.csv | 46 | 0.1398 | 0.354165 | 0.9761 |
| 164 | clim-pet5 | 599.csv | 327 | 0.0511 | 0.357442 | 0.9761 |
| 165 | At1 CFU2 | 63.csv | 46 | 0.1379 | 0.360844 | 0.9761 |
| 166 | clim-tmin9 | 639.csv | 327 | -0.0505 | 0.362568 | 0.9761 |
| 167 | clim-tmin2 | 632.csv | 324 | -0.05 | 0.369497 | 0.9761 |
| 168 | MRVL0 | 641.csv | 51 | 0.1282 | 0.370012 | 0.9761 |
| 169 | B11 | 270.csv | 58 | 0.1189 | 0.374077 | 0.9761 |
| 170 | MS-mean Total length day003 | 730.csv | 48 | 0.1311 | 0.374455 | 0.9761 |
| 171 | root length day001 | 754.csv | 48 | 0.1311 | 0.374455 | 0.9761 |
| 172 | root length day003 | 756.csv | 48 | 0.131 | 0.374761 | 0.9761 |
| 173 | MS-mean Total length day005 | 732.csv | 48 | 0.131 | 0.374761 | 0.9761 |
| 174 | Metal ring in trichomes | 1904.csv | 119 | 0.082 | 0.375518 | 0.9761 |
| 175 | Co59 | 279.csv | 58 | -0.1183 | 0.376469 | 0.9761 |
| 176 | greening 21d protocol a | 1287.csv | 36 | -0.1517 | 0.377265 | 0.9761 |
| 177 | At2 | 83.csv | 46 | 0.133 | 0.378396 | 0.9761 |
| 178 | Mean(LRL) CK | 380.csv | 31 | 0.1635 | 0.379396 | 0.9761 |
| 179 | 0W | 37.csv | 40 | -0.1426 | 0.380205 | 0.9761 |
| 180 | PF-mean Total length day004 | 740.csv | 48 | 0.1283 | 0.384756 | 0.9761 |
| 181 | Yield sweden 2009 (1st experiment) | 139.csv | 68 | -0.1058 | 0.390726 | 0.9761 |
| 182 | 0W GH LN | 82.csv | 33 | -0.1539 | 0.392395 | 0.9761 |
| 183 | root length day007 | 760.csv | 48 | 0.1253 | 0.396024 | 0.9761 |
| 184 | Mean(R) ABA | 370.csv | 30 | -0.1607 | 0.396265 | 0.9761 |
| 185 | Rb85 | 285.csv | 58 | -0.1132 | 0.397584 | 0.9761 |
| 186 | SizeMainEffect2009 | 131.csv | 82 | -0.0943 | 0.399586 | 0.9761 |
| 187 | PF-mean Total length day005 | 741.csv | 48 | 0.1244 | 0.399588 | 0.9761 |
| 188 | callus score 21d protocol a | 1285.csv | 36 | -0.1444 | 0.400876 | 0.9761 |
| 189 | Size spain 2009 (1st experiment) | 128.csv | 82 | -0.0927 | 0.407426 | 0.9761 |
| 190 | Yield spain 2009 (1st experiment) | 124.csv | 63 | 0.1059 | 0.40893 | 0.9761 |
| 191 | MRL0 | 640.csv | 51 | 0.118 | 0.409353 | 0.9761 |
| 192 | Na23 | 271.csv | 58 | -0.1102 | 0.410307 | 0.9761 |
| 193 | shoots 21d protocol b | 1288.csv | 41 | -0.1318 | 0.411448 | 0.9761 |
| 194 | Apical125 | 690.csv | 51 | -0.1167 | 0.41473 | 0.9761 |
| 195 | 8W GH FT | 99.csv | 45 | 0.1244 | 0.415644 | 0.9761 |
| 196 | GR63 cold | 326.csv | 35 | -0.1418 | 0.416352 | 0.9761 |
| 197 | clim-tmin11 | 630.csv | 327 | -0.0442 | 0.425439 | 0.9761 |
| 198 | clim-tmax2 | 620.csv | 327 | -0.0442 | 0.426032 | 0.9761 |
| 199 | root length day005 | 758.csv | 48 | 0.1171 | 0.427976 | 0.9761 |
| 200 | clim-aet9 | 553.csv | 327 | -0.0439 | 0.428538 | 0.9761 |
| 201 | Mean(LRD P) ABA | 374.csv | 30 | -0.1485 | 0.433459 | 0.9761 |
| 202 | clim-gs9 | 587.csv | 327 | -0.0431 | 0.437107 | 0.9761 |
| 203 | clim-aet6 | 550.csv | 327 | -0.043 | 0.438115 | 0.9761 |
| 204 | Shaveproof trichomes | 1909.csv | 120 | 0.0705 | 0.443901 | 0.9761 |
| 205 | greening 21d protocol b | 1294.csv | 41 | -0.1224 | 0.445885 | 0.9761 |

| **Table S13 Screen of the proliferation module eigengene against the AraPheno phenotype collection.** | | | | | | |
|---|---|---|---|---|---|---|
| 206 | clim-tmax4 | 622.csv | 327 | 0.0422 | 0.446698 | 0.9761 |
| 207 | Leaf serr 10 | 6.csv | 46 | -0.1149 | 0.447233 | 0.9761 |
| 208 | Chlorosis 16 | 57.csv | 45 | 0.116 | 0.447806 | 0.9761 |
| 209 | RP GH | 56.csv | 37 | -0.1272 | 0.453232 | 0.9761 |
| 210 | Straight0 | 644.csv | 51 | 0.1074 | 0.453369 | 0.9761 |
| 211 | Mean(TRL) CK | 386.csv | 31 | 0.1394 | 0.45442 | 0.9761 |
| 212 | rFitness tlp | 525.csv | 337 | 0.0408 | 0.455168 | 0.9761 |
| 213 | area 21d protocol a | 1286.csv | 36 | -0.1284 | 0.45552 | 0.9761 |
| 214 | Anthocyanin 16 | 61.csv | 45 | -0.114 | 0.456024 | 0.9761 |
| 215 | Straight75 | 664.csv | 51 | -0.1061 | 0.458594 | 0.9761 |
| 216 | clim-tmax6 | 624.csv | 327 | 0.0409 | 0.461022 | 0.9761 |
| 217 | clim-bio15 | 560.csv | 327 | 0.0409 | 0.461422 | 0.9761 |
| 218 | Mg25 | 272.csv | 58 | 0.0983 | 0.46282 | 0.9761 |
| 219 | FT Field | 86.csv | 48 | 0.1083 | 0.463723 | 0.9761 |
| 220 | Mean(LRD R) C | 357.csv | 30 | -0.1357 | 0.474601 | 0.9807 |
| 221 | clim-gs10 | 577.csv | 327 | -0.0396 | 0.475106 | 0.9807 |
| 222 | Cd111 | 287.csv | 58 | 0.0949 | 0.478597 | 0.9807 |
| 223 | clim-tmin3 | 633.csv | 320 | -0.0396 | 0.47998 | 0.9807 |
| 224 | Rosette Erect 22 | 12.csv | 46 | -0.1062 | 0.482331 | 0.9807 |
| 225 | Germ 10 | 59.csv | 46 | -0.1059 | 0.483581 | 0.9807 |
| 226 | root-like structures 21d protocol b | 1291.csv | 41 | -0.1115 | 0.487596 | 0.9807 |
| 227 | MS-mean Total length day004 | 731.csv | 47 | 0.1031 | 0.490538 | 0.9807 |
| 228 | root length day002 | 755.csv | 47 | 0.1031 | 0.490538 | 0.9807 |
| 229 | Trichome autofluorescence | 1899.csv | 120 | -0.0627 | 0.496591 | 0.9807 |
| 230 | DTFmainEffect2008 | 126.csv | 59 | -0.0889 | 0.503141 | 0.9807 |
| 231 | PC1sc0 | 656.csv | 51 | -0.0958 | 0.503558 | 0.9807 |
| 232 | Mean(LR.no) ABA | 372.csv | 30 | -0.1269 | 0.503998 | 0.9807 |
| 233 | LRDpBZ0 | 647.csv | 51 | -0.0954 | 0.505317 | 0.9807 |
| 234 | Mean(LRD P) CK | 383.csv | 31 | -0.1241 | 0.505874 | 0.9807 |
| 235 | area 15d protocol a | 1272.csv | 36 | -0.1143 | 0.507008 | 0.9807 |
| 236 | DTF spain 2008 (1st experiment) | 132.csv | 59 | -0.0878 | 0.508426 | 0.9807 |
| 237 | RGR P | 719.csv | 48 | -0.0971 | 0.511375 | 0.9807 |
| 238 | LRLpMRL0 | 654.csv | 51 | 0.0938 | 0.512666 | 0.9807 |
| 239 | GR105 cold | 325.csv | 35 | -0.1144 | 0.51304 | 0.9807 |
| 240 | Mean(TLRL) CK | 385.csv | 31 | 0.1213 | 0.515672 | 0.9807 |
| 241 | LC Duration GH | 52.csv | 37 | -0.1099 | 0.517165 | 0.9807 |
| 242 | Inter-specific pollination of Arabidopsis thaliana and Malcolmia littorea | 699.csv | 130 | 0.0571 | 0.518511 | 0.9807 |
| 243 | clim-gs sum | 589.csv | 327 | -0.0358 | 0.518801 | 0.9807 |
| 244 | clim-prec8 | 614.csv | 327 | -0.0352 | 0.526357 | 0.9807 |
| 245 | GR21 warm | 315.csv | 35 | -0.1096 | 0.530653 | 0.9807 |
| 246 | Leaf serr 22 | 92.csv | 46 | 0.0921 | 0.542892 | 0.9807 |
| 247 | aLRL0 | 649.csv | 51 | 0.087 | 0.544016 | 0.9807 |
| 248 | rSurvival fruit tlp | 533.csv | 337 | 0.0332 | 0.544202 | 0.9807 |
| 249 | clim-aet1 | 542.csv | 327 | -0.0334 | 0.546991 | 0.9807 |

| **Table S13 Screen of the proliferation module eigengene against the AraPheno phenotype collection.** | | | | | | |
|---|---|---|---|---|---|---|
| 250 | RGE Fe norm | 723.csv | 48 | 0.0888 | 0.548466 | 0.9807 |
| 251 | root-like structures 15d protocol b | 1277.csv | 41 | -0.096 | 0.550498 | 0.9807 |
| 252 | Plastoquinone | 1314.csv | 35 | -0.1043 | 0.550898 | 0.9807 |
| 253 | callus score 15d protocol b | 1278.csv | 41 | -0.0952 | 0.554007 | 0.9807 |
| 254 | clim-cyclevsgs | 575.csv | 327 | -0.0327 | 0.55623 | 0.9807 |
| 255 | clim-tmax7 | 625.csv | 327 | 0.0324 | 0.558819 | 0.9807 |
| 256 | DTFplantingLoc2008 | 120.csv | 59 | 0.0773 | 0.560418 | 0.9807 |
| 257 | F-mean Total length day003 | 733.csv | 47 | 0.0868 | 0.561665 | 0.9807 |
| 258 | clim-aet8 | 552.csv | 327 | -0.0318 | 0.567006 | 0.9807 |
| 259 | clim-prec9 | 615.csv | 327 | -0.0315 | 0.569793 | 0.9807 |
| 260 | RBN | 706.csv | 453 | -0.0266 | 0.571913 | 0.9807 |
| 261 | Meristem zone length | 143.csv | 37 | 0.0958 | 0.572628 | 0.9807 |
| 262 | PC3sc125 | 698.csv | 51 | -0.0808 | 0.572862 | 0.9807 |
| 263 | GR105 warm | 318.csv | 35 | 0.0985 | 0.573617 | 0.9807 |
| 264 | PC2sc0 | 657.csv | 51 | -0.0806 | 0.573767 | 0.9807 |
| 265 | clim-bio5 | 568.csv | 327 | 0.0311 | 0.575051 | 0.9807 |
| 266 | aLRLpMRL125 | 695.csv | 51 | -0.0802 | 0.575779 | 0.9807 |
| 267 | Autofluorescence color | 1901.csv | 118 | -0.052 | 0.575948 | 0.9807 |
| 268 | As75 | 283.csv | 58 | 0.0733 | 0.584722 | 0.9807 |
| 269 | clim-gs5 | 583.csv | 327 | -0.0303 | 0.585545 | 0.9807 |
| 270 | rFitness mli | 520.csv | 337 | 0.0295 | 0.589395 | 0.9807 |
| 271 | LDV | 34.csv | 44 | -0.0835 | 0.590121 | 0.9807 |
| 272 | clim-pet7 | 601.csv | 327 | 0.0296 | 0.593832 | 0.9807 |
| 273 | RGR Zn norm | 726.csv | 48 | 0.0789 | 0.593833 | 0.9807 |
| 274 | FRI | 69.csv | 42 | -0.0834 | 0.599733 | 0.9807 |
| 275 | Sterol | 1316.csv | 34 | -0.0916 | 0.606585 | 0.9807 |
| 276 | Mean(TRL) C | 359.csv | 30 | 0.0979 | 0.606677 | 0.9807 |
| 277 | clim-aet5 | 549.csv | 327 | -0.0282 | 0.6111 | 0.9807 |
| 278 | LFS GH | 58.csv | 38 | -0.084 | 0.615954 | 0.9807 |
| 279 | clim-bio1 | 554.csv | 327 | -0.0277 | 0.617986 | 0.9807 |
| 280 | clim-pet11 | 594.csv | 327 | -0.0275 | 0.620564 | 0.9807 |
| 281 | clim-bio17 | 562.csv | 327 | -0.0272 | 0.623781 | 0.9807 |
| 282 | Branched0 | 652.csv | 51 | 0.07 | 0.625358 | 0.9807 |
| 283 | undefined structures 15d protocol a | 1269.csv | 36 | -0.0842 | 0.625448 | 0.9807 |
| 284 | MRL125 | 680.csv | 51 | -0.0698 | 0.626418 | 0.9807 |
| 285 | clim-tmax10 | 617.csv | 327 | -0.027 | 0.626526 | 0.9807 |
| 286 | clim-pet1 | 592.csv | 327 | -0.0269 | 0.6283 | 0.9807 |
| 287 | MRVL125 | 681.csv | 51 | -0.069 | 0.630236 | 0.9807 |
| 288 | Metal surrounding trichomes | 1907.csv | 119 | -0.0445 | 0.630804 | 0.9807 |
| 289 | MRVA125 | 683.csv | 51 | 0.0687 | 0.63208 | 0.9807 |
| 290 | undefined structures 21d protocol a | 1283.csv | 36 | -0.0823 | 0.633239 | 0.9807 |
| 291 | clim-aet2 | 546.csv | 327 | -0.0264 | 0.633801 | 0.9807 |
| 292 | K39 | 275.csv | 58 | -0.0629 | 0.639225 | 0.9807 |
| 293 | aLRL125 | 689.csv | 51 | -0.0669 | 0.640668 | 0.9807 |
| 294 | RGR P norm | 724.csv | 48 | -0.0689 | 0.641682 | 0.9807 |

| **Table S13 Screen of the proliferation module eigengene against the AraPheno phenotype collection.** | | | | | | |
|---|---|---|---|---|---|---|
| 295 | MT GH | 38.csv | 37 | -0.0787 | 0.643385 | 0.9807 |
| 296 | Leaf serr 16 | 3.csv | 45 | 0.0709 | 0.643449 | 0.9807 |
| 297 | noLRpMR0 | 645.csv | 51 | 0.0661 | 0.644783 | 0.9807 |
| 298 | clim-bio3 | 566.csv | 327 | -0.025 | 0.652044 | 0.9807 |
| 299 | clim-prec11 | 606.csv | 327 | -0.0246 | 0.657756 | 0.9807 |
| 300 | clim-bio14 | 559.csv | 327 | -0.0245 | 0.659539 | 0.9807 |
| 301 | FT Duration GH | 32.csv | 37 | -0.0743 | 0.662152 | 0.9807 |
| 302 | clim-pet4 | 598.csv | 327 | 0.0242 | 0.662918 | 0.9807 |
| 303 | clim-prec3 | 609.csv | 327 | 0.0241 | 0.664147 | 0.9807 |
| 304 | root length day004 minusZn | 764.csv | 48 | 0.0643 | 0.664302 | 0.9807 |
| 305 | 8W GH LN | 36.csv | 45 | 0.0664 | 0.664801 | 0.9807 |
| 306 | PC2sc75 | 677.csv | 51 | 0.0619 | 0.666254 | 0.9807 |
| 307 | shoot primordia 15d protocol b | 1275.csv | 41 | -0.069 | 0.668138 | 0.9807 |
| 308 | clim-tmax8 | 626.csv | 327 | 0.0237 | 0.669242 | 0.9807 |
| 309 | MRVL75 | 661.csv | 51 | -0.0607 | 0.672418 | 0.9807 |
| 310 | RGR P Zn | 720.csv | 48 | -0.0611 | 0.680085 | 0.9807 |
| 311 | Mean(P2) CK | 382.csv | 31 | 0.0766 | 0.681936 | 0.9807 |
| 312 | YieldPlantingSummerLocSweden2009 | 138.csv | 60 | 0.0536 | 0.684179 | 0.9807 |
| 313 | callus score 15d protocol a | 1271.csv | 36 | -0.0698 | 0.685852 | 0.9807 |
| 314 | Chlorosis 10 | 55.csv | 46 | 0.0612 | 0.686117 | 0.9807 |
| 315 | clim-aet3 | 547.csv | 327 | -0.022 | 0.691791 | 0.9807 |
| 316 | rFitness mlp | 521.csv | 337 | 0.0216 | 0.69247 | 0.9807 |
| 317 | Mean(TLRL) C | 358.csv | 30 | -0.0735 | 0.699407 | 0.9807 |
| 318 | Depth75 | 662.csv | 51 | -0.0552 | 0.700574 | 0.9807 |
| 319 | noLRpMR75 | 665.csv | 51 | -0.0545 | 0.703854 | 0.9807 |
| 320 | clim-prec1 | 604.csv | 327 | 0.0207 | 0.709016 | 0.9807 |
| 321 | Yield spain 2009 (2nd experiment) | 122.csv | 55 | 0.0511 | 0.710982 | 0.9807 |
| 322 | Mean(LRD R) CK | 384.csv | 31 | -0.0675 | 0.718165 | 0.9807 |
| 323 | clim-pet6 | 600.csv | 327 | 0.02 | 0.718306 | 0.9807 |
| 324 | clim-aet7 | 551.csv | 327 | -0.0199 | 0.719902 | 0.9807 |
| 325 | Mean(LRR) CK | 387.csv | 31 | 0.0667 | 0.721511 | 0.9807 |
| 326 | Mn55 | 277.csv | 58 | 0.0475 | 0.723388 | 0.9807 |
| 327 | clim-gs prec | 588.csv | 327 | -0.0195 | 0.724857 | 0.9807 |
| 328 | PC3sc75 | 678.csv | 51 | -0.0499 | 0.728234 | 0.9807 |
| 329 | Depth125 | 682.csv | 51 | -0.0496 | 0.729469 | 0.9807 |
| 330 | Mature cell length | 142.csv | 37 | 0.0588 | 0.729709 | 0.9807 |
| 331 | YieldMainEffect2009 | 137.csv | 80 | 0.0391 | 0.73042 | 0.9807 |
| 332 | clim-gs6 | 584.csv | 327 | -0.0189 | 0.733512 | 0.9807 |
| 333 | Fe57 | 278.csv | 58 | 0.0457 | 0.733571 | 0.9807 |
| 334 | Diameter | 707.csv | 435 | -0.0162 | 0.73554 | 0.9807 |
| 335 | LRDpBZ125 | 687.csv | 51 | -0.0481 | 0.737601 | 0.9807 |
| 336 | MRL75 | 660.csv | 51 | -0.0474 | 0.741248 | 0.9807 |
| 337 | PZ-mean Total length day003 | 742.csv | 47 | 0.0493 | 0.742345 | 0.9807 |
| 338 | clim-prec2 | 608.csv | 327 | 0.0182 | 0.742536 | 0.9807 |
| 339 | clim-aet12 | 545.csv | 327 | -0.0181 | 0.744922 | 0.9807 |

| **Table S13 Screen of the proliferation module eigengene against the AraPheno phenotype collection.** | | | | | | |
|---|---|---|---|---|---|---|
| 340 | shoots 21d protocol a | 1281.csv | 36 | -0.0556 | 0.747474 | 0.9807 |
| 341 | Mean(LR.no) CK | 381.csv | 31 | -0.0599 | 0.748938 | 0.9807 |
| 342 | RGR P Fe norm | 727.csv | 48 | 0.0471 | 0.75036 | 0.9807 |
| 343 | Mean(TRL) ABA | 377.csv | 30 | 0.0604 | 0.751256 | 0.9807 |
| 344 | rSeeds mhp | 535.csv | 337 | 0.0173 | 0.75162 | 0.9807 |
| 345 | rSurvival fruit thi | 530.csv | 337 | 0.0173 | 0.75164 | 0.9807 |
| 346 | clim-gs2 | 580.csv | 327 | 0.0172 | 0.756024 | 0.9807 |
| 347 | Ca43 | 276.csv | 58 | 0.0416 | 0.756559 | 0.9807 |
| 348 | P-mean Total length day004 | 737.csv | 48 | -0.0457 | 0.757757 | 0.9807 |
| 349 | clim-tmax3 | 621.csv | 327 | -0.0171 | 0.757834 | 0.9807 |
| 350 | Mo98 | 286.csv | 58 | -0.0406 | 0.762074 | 0.9807 |
| 351 | clim-bio10 | 555.csv | 327 | 0.0167 | 0.763984 | 0.9807 |
| 352 | RGR P Fe | 722.csv | 48 | 0.0444 | 0.764292 | 0.9807 |
| 353 | Mean(R) CK | 379.csv | 31 | -0.0561 | 0.764524 | 0.9807 |
| 354 | clim-gs11 | 578.csv | 327 | -0.0166 | 0.764925 | 0.9807 |
| 355 | LRL125 | 688.csv | 51 | 0.0429 | 0.765186 | 0.9807 |
| 356 | undefined structures 15d protocol b | 1276.csv | 41 | -0.0479 | 0.766003 | 0.9807 |
| 357 | MRLpTRS125 | 693.csv | 51 | -0.0421 | 0.769075 | 0.9807 |
| 358 | shoot primordia 15d protocol a | 1268.csv | 36 | -0.0506 | 0.76925 | 0.9807 |
| 359 | clim-bio19 | 564.csv | 327 | 0.0161 | 0.771956 | 0.9807 |
| 360 | clim-pet2 | 596.csv | 327 | -0.016 | 0.772725 | 0.9807 |
| 361 | Basal0 | 651.csv | 51 | -0.0409 | 0.775472 | 0.9807 |
| 362 | clim-bio18 | 563.csv | 327 | -0.0153 | 0.782423 | 0.9807 |
| 363 | clim-bio9 | 572.csv | 326 | -0.0153 | 0.782763 | 0.9807 |
| 364 | rSeeds mlp | 537.csv | 126 | -0.0245 | 0.7851 | 0.9807 |
| 365 | clim-bio16 | 561.csv | 327 | 0.015 | 0.787323 | 0.9807 |
| 366 | MRVA75 | 663.csv | 51 | -0.0384 | 0.789211 | 0.9807 |
| 367 | Apical0 | 650.csv | 51 | 0.0373 | 0.794941 | 0.9807 |
| 368 | Basal75 | 671.csv | 51 | 0.036 | 0.802173 | 0.9807 |
| 369 | PC1sc75 | 676.csv | 51 | 0.0357 | 0.803466 | 0.9807 |
| 370 | shoot primordia 21d protocol a | 1282.csv | 36 | -0.0424 | 0.80608 | 0.9807 |
| 371 | clim-prec6 | 612.csv | 327 | 0.0134 | 0.808949 | 0.9807 |
| 372 | clim-cyclelength | 573.csv | 327 | -0.0134 | 0.80904 | 0.9807 |
| 373 | Root length (Day 3) | 291.csv | 33 | 0.0437 | 0.809048 | 0.9807 |
| 374 | P-mean Total length day005 | 738.csv | 48 | -0.0357 | 0.809545 | 0.9807 |
| 375 | clim-pet12 | 595.csv | 327 | -0.0126 | 0.82053 | 0.9807 |
| 376 | Root length (Day 1) | 289.csv | 32 | -0.0413 | 0.822308 | 0.9807 |
| 377 | As CFU2 | 70.csv | 46 | 0.0332 | 0.826597 | 0.9807 |
| 378 | Width 10 | 18.csv | 46 | -0.0328 | 0.828434 | 0.9807 |
| 379 | clim-gs3 | 581.csv | 327 | 0.012 | 0.828446 | 0.9807 |
| 380 | Li7 | 269.csv | 58 | -0.029 | 0.828831 | 0.9807 |
| 381 | aLRLpMRL75 | 675.csv | 51 | -0.031 | 0.82897 | 0.9807 |
| 382 | PZ-mean Total length day004 | 743.csv | 48 | 0.0317 | 0.830582 | 0.9807 |
| 383 | stomatasize | 750.csv | 190 | -0.0156 | 0.830602 | 0.9807 |
| 384 | Leaf roll 16 | 8.csv | 45 | 0.0326 | 0.831804 | 0.9807 |

| **Table S13 Screen of the proliferation module eigengene against the AraPheno phenotype collection.** | | | | | | |
|---|---|---|---|---|---|---|
| 385 | clim-pet10 | 593.csv | 327 | -0.0118 | 0.832072 | 0.9807 |
| 386 | clim-tmin8 | 638.csv | 327 | -0.0116 | 0.834435 | 0.9807 |
| 387 | clim-prec10 | 605.csv | 327 | -0.0116 | 0.834675 | 0.9807 |
| 388 | aLRLpMRL0 | 655.csv | 51 | 0.0299 | 0.834899 | 0.9807 |
| 389 | RGR P Zn norm | 725.csv | 48 | -0.0305 | 0.837116 | 0.9807 |
| 390 | clim-pet8 | 602.csv | 327 | 0.0113 | 0.839016 | 0.9807 |
| 391 | Metals in stomata | 1908.csv | 119 | 0.0186 | 0.841051 | 0.9807 |
| 392 | Leaf roll 22 | 46.csv | 46 | -0.0301 | 0.842543 | 0.9807 |
| 393 | aLRL75 | 669.csv | 51 | 0.0284 | 0.843231 | 0.9807 |
| 394 | clim-aet4 | 548.csv | 327 | -0.0108 | 0.845919 | 0.9807 |
| 395 | Width 16 | 14.csv | 45 | -0.0296 | 0.846707 | 0.9807 |
| 396 | PC2sc125 | 697.csv | 51 | 0.0274 | 0.848868 | 0.9807 |
| 397 | undefined structures 21d protocol b | 1290.csv | 41 | 0.0305 | 0.849827 | 0.9807 |
| 398 | LRLpMRL1225 | 694.csv | 51 | 0.0269 | 0.85157 | 0.9807 |
| 399 | Mean(LRL) C | 353.csv | 30 | -0.0354 | 0.852837 | 0.9807 |
| 400 | LRDpMR0 | 646.csv | 51 | -0.0266 | 0.852896 | 0.9807 |
| 401 | Root length (Day 2) | 290.csv | 33 | 0.0335 | 0.853018 | 0.9807 |
| 402 | clim-pet9 | 603.csv | 327 | 0.0101 | 0.856002 | 0.9808 |
| 403 | clim-bio13 | 558.csv | 327 | 0.01 | 0.857429 | 0.9808 |
| 404 | LRDpMR75 | 666.csv | 51 | -0.0244 | 0.865158 | 0.9832 |
| 405 | clim-gs7 | 585.csv | 327 | -0.0093 | 0.867345 | 0.9832 |
| 406 | Anthocyanin 10 | 62.csv | 46 | 0.0247 | 0.870712 | 0.9832 |
| 407 | PC3sc0 | 658.csv | 51 | -0.0233 | 0.87123 | 0.9832 |
| 408 | clim-tmax9 | 627.csv | 327 | 0.0089 | 0.872244 | 0.9832 |
| 409 | clim-bio12 | 557.csv | 327 | -0.0087 | 0.876035 | 0.9832 |
| 410 | MRLpTRS75 | 673.csv | 51 | 0.0222 | 0.877357 | 0.9832 |
| 411 | clim-prec5 | 611.csv | 327 | -0.0083 | 0.880835 | 0.9832 |
| 412 | clim-gs tmax | 590.csv | 327 | -0.0083 | 0.881223 | 0.9832 |
| 413 | LRLpMRL75 | 674.csv | 51 | 0.0208 | 0.884667 | 0.9832 |
| 414 | Mean(LRL) ABA | 371.csv | 30 | -0.0276 | 0.884867 | 0.9832 |
| 415 | Mean(R) C | 352.csv | 30 | -0.027 | 0.887565 | 0.9832 |
| 416 | clim-pet3 | 597.csv | 327 | -0.0077 | 0.889133 | 0.9832 |
| 417 | Stomatasize 261 | 751.csv | 149 | -0.0115 | 0.889395 | 0.9832 |
| 418 | clim-gs8 | 586.csv | 327 | 0.0074 | 0.893926 | 0.9859 |
| 419 | clim-prec12 | 607.csv | 327 | 0.007 | 0.899303 | 0.9869 |
| 420 | PZ-mean Total length day005 | 744.csv | 48 | 0.0185 | 0.900865 | 0.9869 |
| 421 | clim-gs4 | 582.csv | 327 | 0.0067 | 0.904439 | 0.9869 |
| 422 | Root length (Day 4) | 292.csv | 30 | 0.0225 | 0.906096 | 0.9869 |
| 423 | Metal in trichome base | 1906.csv | 119 | 0.0108 | 0.907302 | 0.9869 |
| 424 | LRL75 | 668.csv | 51 | 0.0166 | 0.907683 | 0.9869 |
| 425 | clim-gs12 | 579.csv | 327 | -0.006 | 0.913898 | 0.988 |
| 426 | root length day006 minusZn | 766.csv | 48 | -0.0157 | 0.915874 | 0.988 |
| 427 | MRLpTRS0 | 653.csv | 51 | -0.0151 | 0.916416 | 0.988 |
| 428 | GR21 | 294.csv | 57 | 0.0139 | 0.918074 | 0.988 |
| 429 | P31 | 273.csv | 58 | 0.0129 | 0.923658 | 0.988 |

| **Table S13 Screen of the proliferation module eigengene against the AraPheno phenotype collection.** | | | | | | |
|---|---|---|---|---|---|---|
| 430 | 8W | 48.csv | 45 | -0.0147 | 0.923773 | 0.988 |
| 431 | noLRpMR125 | 685.csv | 51 | -0.0134 | 0.925492 | 0.988 |
| 432 | Bs CFU2 | 42.csv | 46 | -0.0141 | 0.925838 | 0.988 |
| 433 | clim-tmin5 | 635.csv | 327 | 0.0045 | 0.935645 | 0.9882 |
| 434 | Ni60 | 280.csv | 58 | -0.0108 | 0.935824 | 0.9882 |
| 435 | Tokoferol | 1317.csv | 35 | 0.0141 | 0.935878 | 0.9882 |
| 436 | deltaRGR P Fe norm on Fe norm | 728.csv | 48 | -0.0118 | 0.936401 | 0.9882 |
| 437 | rSeeds mli | 536.csv | 282 | 0.0047 | 0.937549 | 0.9882 |
| 438 | rSurvival fruit mhp | 527.csv | 337 | 0.004 | 0.942042 | 0.9882 |
| 439 | Area sweden 2009 (2nd experiment) | 112.csv | 83 | 0.008 | 0.943064 | 0.9882 |
| 440 | clim-tmin6 | 636.csv | 327 | 0.0038 | 0.944739 | 0.9882 |
| 441 | Chlorosis 22 | 19.csv | 46 | 0.0104 | 0.945329 | 0.9882 |
| 442 | Branched75 | 672.csv | 51 | 0.0095 | 0.947428 | 0.9882 |
| 443 | Branched125 | 692.csv | 51 | -0.0089 | 0.950779 | 0.9891 |
| 444 | clim-prec7 | 613.csv | 327 | -0.0033 | 0.952629 | 0.9891 |
| 445 | Straight125 | 684.csv | 51 | 0.0075 | 0.958143 | 0.9907 |
| 446 | PC1sc125 | 696.csv | 51 | -0.0075 | 0.958423 | 0.9907 |
| 447 | shoot primordia 21d protocol b | 1289.csv | 41 | 0.0075 | 0.962957 | 0.9931 |
| 448 | TRS125 | 679.csv | 51 | 0.0053 | 0.97074 | 0.9978 |
| 449 | Polyprenol | 1315.csv | 34 | -0.0048 | 0.978629 | 0.9978 |
| 450 | P-mean Total length day003 | 736.csv | 45 | 0.004 | 0.978939 | 0.9978 |
| 451 | LRDpMR125 | 686.csv | 51 | 0.003 | 0.983078 | 0.9978 |
| 452 | clim-tmin7 | 637.csv | 327 | 0.0009 | 0.987555 | 0.9978 |
| 453 | callus score 21d protocol b | 1292.csv | 41 | -0.0023 | 0.988495 | 0.9978 |
| 454 | Zn66 | 282.csv | 58 | -0.0017 | 0.989745 | 0.9978 |
| 455 | root-like structures 15d protocol a | 1270.csv | 36 | -0.002 | 0.990534 | 0.9978 |
| 456 | GR63 warm | 320.csv | 35 | 0.0021 | 0.990664 | 0.9978 |
| 457 | TRS75 | 659.csv | 51 | 0.0016 | 0.990874 | 0.9978 |
| 458 | clim-tmin4 | 634.csv | 327 | 0.0004 | 0.993777 | 0.9978 |
| 459 | root-like structures 21d protocol a | 1284.csv | 36 | 0.0013 | 0.993914 | 0.9978 |
| 460 | rFitness mhp | 519.csv | 337 | 0.0003 | 0.995626 | 0.9978 |
| 461 | Root length (Day 5) | 293.csv | 32 | -0.0004 | 0.998335 | 0.9983 |
